# Origins, admixture dynamics and homogenization of the African gene pool in the Americas

**DOI:** 10.1101/652701

**Authors:** Mateus H. Gouveia, Victor Borda, Thiago P. Leal, Rennan G. Moreira, Andrew W. Bergen, Marla M. Aquino, Gilderlanio S. Araujo, Nathalia M. Araujo, Fernanda S.G. Kehdy, Raquel Liboredo, Moara Machado, Wagner C.S. Magalhaes, Lucas A. Michelin, Maíra R. Rodrigues, Fernanda Rodrigues-Soares, Hanaisa P. Sant Anna, Meddly L. Santolalla, Marília O. Scliar, Giordano Soares-Souza, Roxana Zamudio, Camila Zolini, Michael Dean, Robert H. Gilman, Heinner Guio, Jorge Rocha, Alexandre C. Pereira, Mauricio L. Barreto, Bernardo L. Horta, Maria F. Lima-Costa, Sam M. Mbulaiteye, Stephen J. Chanock, Sarah A. Tishkoff, Meredith Yeager, Eduardo Tarazona-Santos

## Abstract

The Transatlantic Slave Trade transported more than 9 million Africans to the Americas between the early 16th and the mid-19th centuries. We performed genome-wide analysis of 6,267 individuals from 22 populations and observed an enrichment in West-African ancestry in northern latitudes of the Americas, whereas South/East African ancestry is more prevalent in southern South-America. This pattern results from distinct geographic and geopolitical factors leading to population differentiation. However, we observed a decrease of 68% in the African gene pool between-population diversity within the Americas when compared to the regions of origin from Africa, underscoring the importance of historical factors favoring admixture between individuals with different African origins in the New World. This is consistent with the excess of West-Central Africa ancestry (the most prevalent in the Americas) in the US and Southeast-Brazil, respect to historical-demography expectations. Also, in most of the Americas, admixture intensification occurred between 1,750 and 1,850, which correlates strongly with the peak of arrivals from Africa. This study contributes with a population genetics perspective to the ongoing social, cultural and political debate regarding ancestry, race, and admixture in the Americas.

**Significance Statement:** Differently from most genetic studies, that have estimated the overall African ancestry in the Americas, we perform a finer geographic analysis and infer how different African groups contributed to North-, South-American and Caribbean populations, in the context of geographic and geopolitical factors. We also perform a formal comparison of information from demographic history records of the Transatlantic Slave Trade with inferences based on genomic diversity of current populations. Our approach reveals the distinct regional African ancestry roots of different populations from North-, South-America and the Caribe and other important aspects of the historical process of *mestizaje* and its dynamics in the American continent.

## Introduction

The Transatlantic Slave Trade was an international enterprise involving Brazilian, British, Danish, Dutch, French, German, Portuguese, Spanish and Swedish traders. They brought over 9 million Africans to the Americas between the early 16^th^ and the mid-19^th^ centuries. African regions of origin included far away locations as Senegambia is from Tanzania. Destiny ports in the Americas were also distant as Newport in New England is from Buenos Aires (1, 2). The Transatlantic Slave Trade shaped the genetic structure of American continent populations (3–11). While most genetic studies have estimated the overall African ancestry in the Americas, a finer geographic analysis is needed to infer how different African groups contributed to North-, Central-, South-American and Caribbean populations. The geopolitical factors that permeated the African Diaspora have been seldom discussed at a continental scale, despite its potential influence on the genetic structure of populations. Furthermore, a formal comparison of information from demographic history records of the Transatlantic Slave Trade with inferences based on genomic diversity of current populations has not been performed.

Here we perform a joint analysis of genetic data and historical records of the Transatlantic Slave Trade to address the following questions: (i) Is there a correspondence between the geographic origin of specific African populations of the Diaspora and specific destinations in the Americas?; (ii) Was admixture dynamics in the Americas associated with the dynamics of arrivals of African slaves?; (iii) Considering the geographic extension and the massive demographic magnitude of the African Diaspora, as well as the level of *between-populations* genetic differentiation in the African regions of origin of slaves, did the Transatlantic Slave Trade lead to a higher, similar or lower level of *between-population* differentiation of the African gene pool in the Americas?

## Results and Discussion

We combined genome-wide data from 25 populations: 9 admixed of the Americas, 11 Africans, 2 Europeans and 3 Native Americans (Fig. 1, *SI Appendix*, Table S1 and section 1.1) and created a dataset of 6,267 unrelated individuals with >10% of African ancestry (*SI Appendix*, sections 1 and 2). Using ADMIXTURE (12), we identified two continental (European and Native American) and four African-specific ancestry clusters, named based on their association with geographic regions (*SI Appendix*, Table S1, represented by different colors in Fig. 1): (i) West-Central African (blue), (ii) Western African (purple) and, (iii) South/East African (yellow), which are prevalent in the Americas, as well as (iv) Northern Ugandan (NU, cyan), which accounts for a very low proportion of African ancestry in the Americas. Hereafter, while in African individuals, the proportion of ADMIXTURE ancestry clusters are relative to their whole genome ancestry (Fig. 1A, *SI Appendix*, Table S1), in American continent individuals, these proportions are relative to the sum of the four African ancestry clusters (Fig 1A). We also estimated haplotype-based population admixture proportions (13, 14) in populations of the Americas, that in Fig. 2A and B, and *SI Appendix*, Table S3, are relative to the total contribution of African populations.

**Fig. 1.**
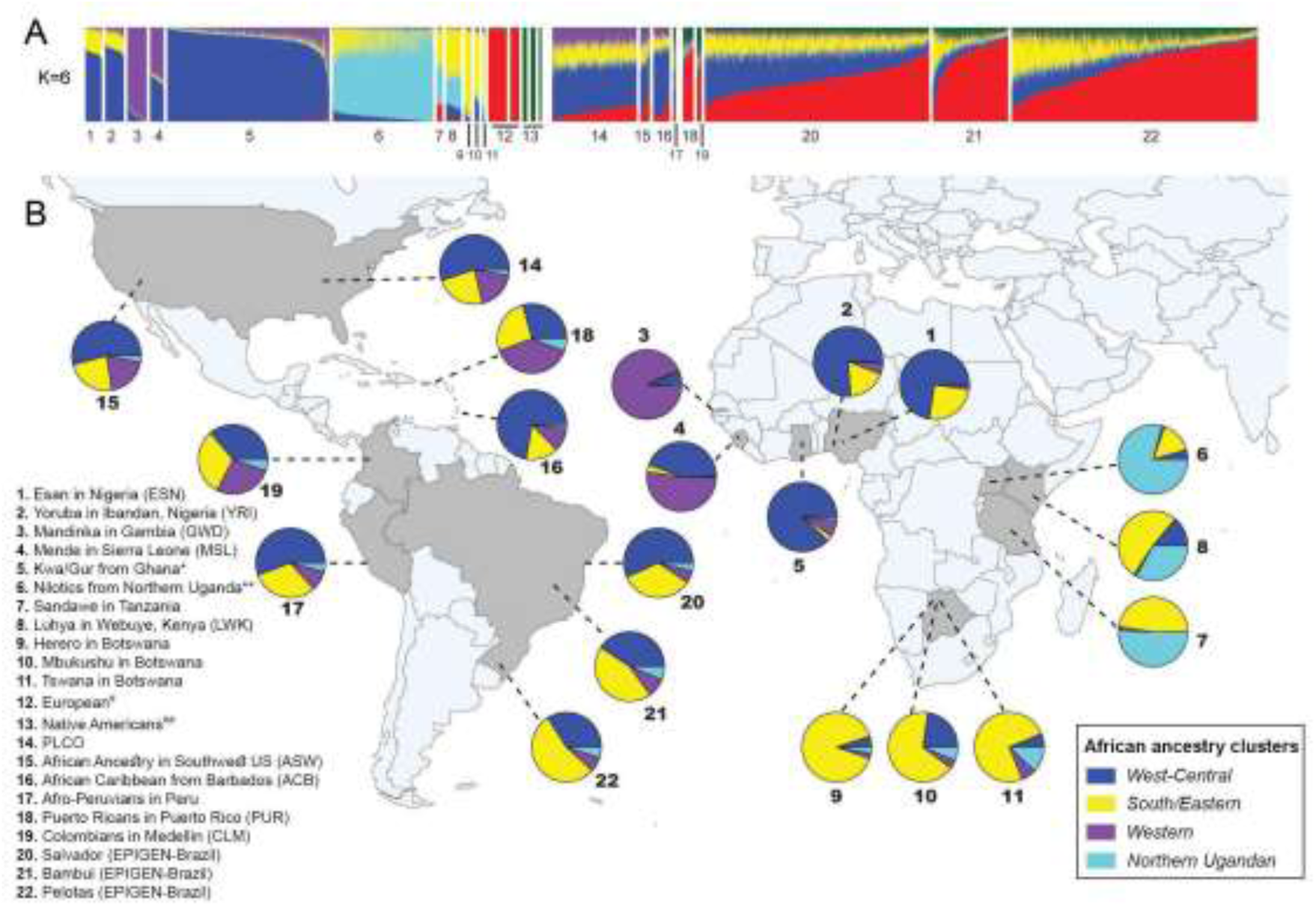
Ancestry analysis of African and admixed populations of the Americas inferred using ADMIXTURE (K=6). (A) Vertical bar plot showing the total African, European and Native American proportions of the ancestry clusters (*SI Appendix*, Fig. S1 and section 2.6.1.). *The Kwa/Gur dataset includes approximately 35 ethno-linguistic groups, predominantly from the Kwa and Gur Niger-Congo linguistic group(27). **The Nilotics dataset includes predominantly three ethno-linguistic groups in Northern Uganda (Langi, Acholi and Lugbara) from the Nilotic linguistic group^27^; ^#^the Europeans are: Iberian Population in Spain (IBS) and Utah residents with Northern and Western European ancestry (CEU), in this order in the ADMIXTURE bar plot; ^##^the Native Americans are: Shimaa, Ashaninka and Aymara, respectively from Borda et al. (in preparation); the PLCO (Prostate, Lung, Colorectal and Ovarian Cancer Screening) data is comprised of African-Americans from East USA. (B) Percentages of subcontinental African ancestry clusters. For admixed populations of the American continent these percentages are relative to the total African ancestry (i.e. the sum of the four African associated clusters: West-Central, Western, Southern/Eastern, Northern Ugandan).

**Fig. 2.**
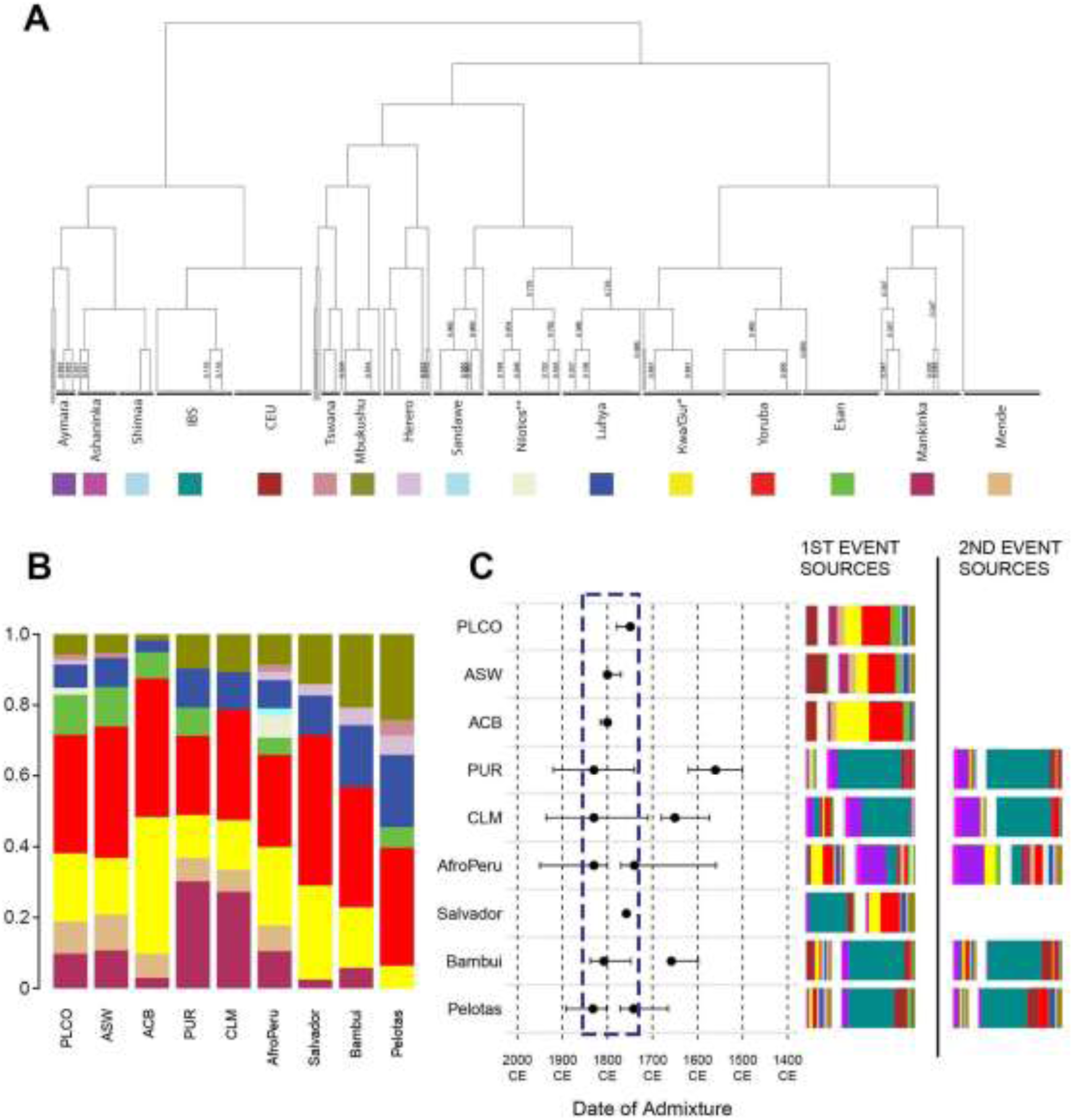
Haplotype-based clustering of parental individuals and admixture inferences for admixed American continent populations. (A) fineSTRUCTURE tree of parental individuals. *The Kwa/Gur dataset includes approximately 35 ethno-linguistic groups, predominantly from the Kwa and Gur linguistic group^27^. **The Nilotics dataset includes predominantly three ethno-linguistic groups from Northern Uganda (Langi, Acholi and Lugbara) of the Nilotic linguistic group^27^. (B) Subcontinental contributions relative to the total African ancestry in admixed populations inferred by the MIXTURE MODEL. (C) GLOBETROTTER inference of admixture events for each admixed population. Inferred date(s) and 95% confidence intervals are represented by dots and horizontal lines in the graph, respectively. Dashed rectangle in the admixture dates plots highlights the most dynamic period for admixture. Beside the dating graph, we represented the inferred admixing sources (bars) for one and two events. Bar size represents the genetic contribution of the source. Each color corresponds to the contribution of each parental population. CEU=Utah Residents (CEPH) with Northern and Western Ancestry-USA, IBS=Iberian population in Spain, CLM=Colombians from Medellin, PUR=Puerto Ricans from Puerto Rico, ACB=African Caribbeans in Barbados, ASW= African Americans in Southwest USA, PLCO= African Americans from East USA.

### Ancestry correspondence between African and admixed American continent populations, and the influence of geography and geopolitics

The West-Central Africa-associated ancestry cluster is the most prevalent African cluster in the Americas, including African-Caribbean from Barbados (72% of African ancestry), Northeastern Brazilians (57%), Afro-Peruvians (56%) and US African-Americans (54-55%) (blue in Fig. 1, *SI Appendix*, Table S1 and section 2.1). Moreover, haplotype-based analysis(13, 14) reveals a higher contribution from Yoruba-like and Esan-like populations (from Nigeria, mean: 38%) than from Kwa/Gur-like populations (from Ghana, mean: 18%) (Fig. 2A and B, *SI Appendix*, Table S3).

The Western Africa-associated ancestry cluster has its highest proportions in Puerto Ricans (38% of African ancestry), Colombians (27%) and US African-Americans (19-20%, purple in Fig. 1, *SI Appendix*, Table S1), while Brazilians have the lowest proportion (<9%), limited to a Mandinka-like (Gambia) contribution and with no Mende-like (Sierra Leone) contribution (Fig. 2A and B, *SI Appendix*, Table S3).

The South/East Africa-associated ancestry cluster, in contrast, shows its highest proportion in South and Southeast Brazil (44% and 54% of total African ancestry, respectively) (yellow in Fig. 1 and *SI Appendix*, Table S1). Haplotype-based methods (13, 14) identified two different sources of gene flow associated with the South/Eastern Africa ancestry cluster: one from Mbukushu-like populations (Botswana, Western Bantu speakers from Southern Africa, 20-24% to South/Southeast Brazil) and one from Luhya-like populations (Kenya, Eastern Bantu speakers from Eastern Africa, 17-20% to South/Southeast Brazil, Fig. 2A and B, *SI Appendix*, Table S3). Western- and Eastern-Bantu speakers historically correspond to the two streams of the Bantu migrations in the last 4000-2500 years (10, 15, 16).

This emerging portrait of the African ancestry in the Americas suggests an influence of geography and geopolitics. Geographical factors include: (i) the latitudinal proximity between Western Africa and Caribe-Central/North America, as well as between South/East Africa and Southern Brazil, (ii) the winds and ocean currents, that shaped two navigation systems: the North-Atlantic, with voyages mostly to North America, and the South-Atlantic, with voyages predominantly to Brazil (17). Indeed, West Central Africa- and West Africa-associated ancestry clusters are more commonly observed in northern latitudes, while the South/East Africa-associated ancestry cluster is more evident in southern latitudes.

Differently, the Portugal possessions in the Americas (Brazil) and its influence in South and East African coasts (current Angola and Mozambique)(18) exemplify the geopolitical factors that affected, in particular, the distribution of the South/East Africa-associated ancestry cluster. While the Portuguese Crown had earlier privileged relations with the kingdoms of Benin in nowadays Nigeria, it later extended its influence to Bantu-speaking areas such as Congo/Angola and Mozambique (19). Indeed, Portuguese-Brazilian slave trade routes departed from Luanda and Cabinda (Angola) and from Zanzibar (Tanzania) and Inhambane (Mozambique) during 18^th^ and 19^th^ centuries (1). The abolition of slavery by the British in 1807, who controlled the North Atlantic route, also led Portuguese traders to prefer routes in the South Atlantic(20). Therefore, geography (inter-continental distances and climatic factors affecting transatlantic navigation) and geopolitics (European colonial influences and possessions) influenced the geographic and linguistic diversity of African emigrants as well as favored the regional differentiation of African ancestry in the Americas.

### The dynamics of African admixture in the Americas accompanied the dynamics of arrivals of African slaves

Remarkably, linkage-disequilibrium-based inference (14) shows that all the studied admixed populations of the Americas exhibit the signature of an intensification of admixture in the interval from 1750 to 1850 (Fig. 2C, *SI Appendix*, section 3), revealing a continental trend. This trend is consistent with results by Baharian et al. (3), focused in the US. Importantly, this time interval matches or is immediately subsequent to regional peaks of number of slaves arriving from Africa to US, Barbados, Puerto Rico and Brazil (*SI Appendix*, Fig. S4). Thus, in most of the Americas, the arrival of the largest contingent of Africans between 1700 and 1850 (*SI Appendix*, Figs. S4) was almost synchronic with intensive admixture, a process that was also characterized by positive ancestry-based assortative mating (6).

### The African gene pool is more homogenous between-populations in the Americas than in Africa

Figure 1 suggests that African ancestry clusters are more homogeneously distributed between admixed American continent populations than between the African populations that contributed to the Transatlantic Slave Trade. Considering only the African gene pool, the largest differentiation, measured by the *African-Specific Genetic Distance* (*ASGD*, Fig. 3 and *SI Appendix*, section 4 and Figs. S5), is observed between African populations (mean: 0.057, mean excluding populations with marginal contribution to the Americas [Nilotics and Sandawe: 0.53]), followed by differentiation between African vs. America’s populations (mean: 0.043) and between populations of the Americas (mean: 0.018, 32% of the ASGD between African populations) (Wilcoxon test, p < 10^−6^ for the three pairwise comparisons, Fig. 3).

**Fig. 3.**
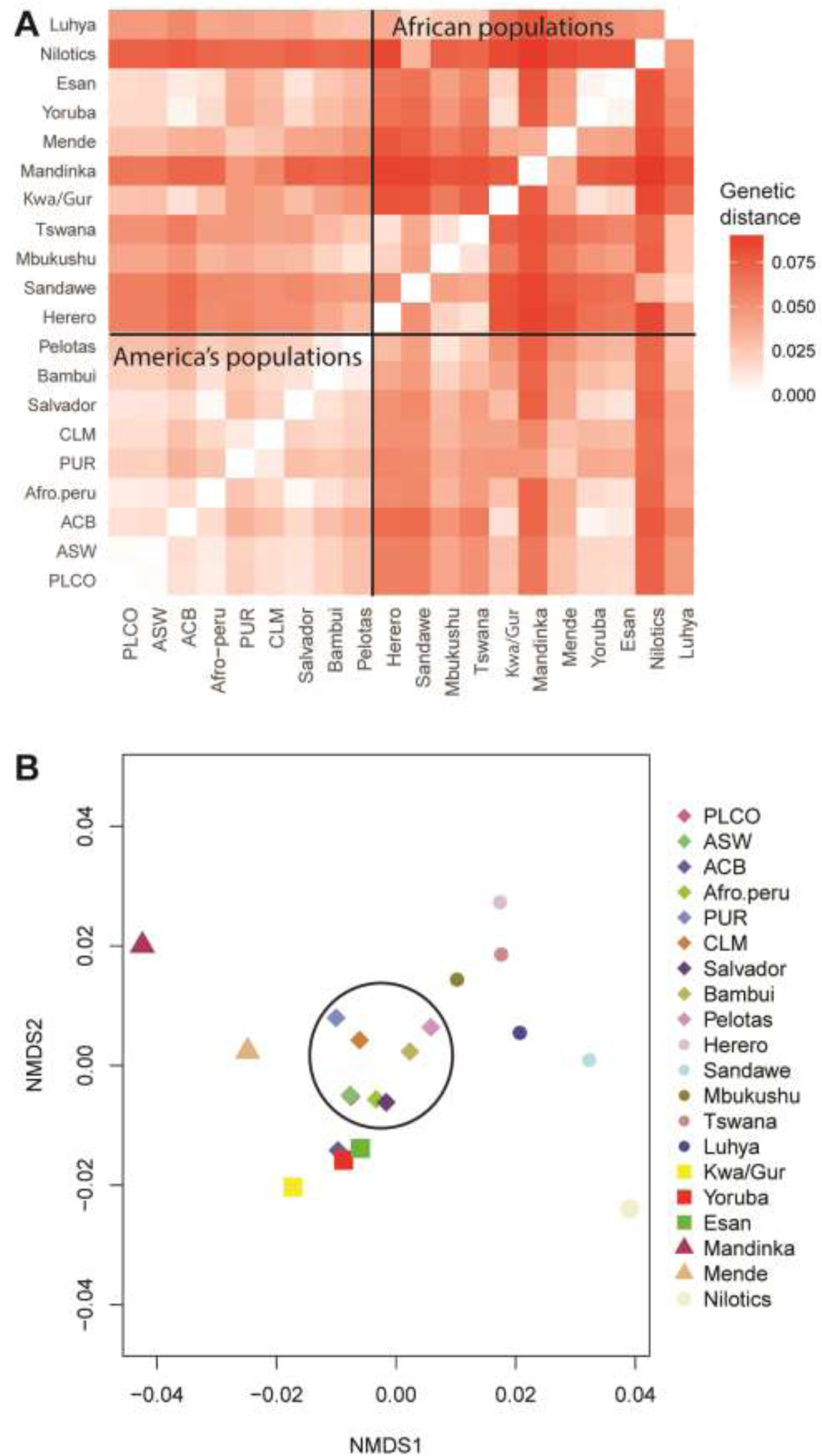
Pairwise genetic distances of the African gene pool between populations of the American continent and Africa. (A) Heatmap Matrix and (B) Multidimensional scaling of the African gene pool genetic distances. We used solid squares, triangles and circles to represent populations associate with WCA: West-Central Africa, SEA: South/East Africa and WA: Western Africa ancestry clusters. The admixed American continent populations are highlighted with the circle with the exception of the ACB population which clustered closer to the WCA ancestry-associated populations. CLM=Colombians from Medellin, PUR=Puerto Ricans from Puerto Rico, ACB=African Caribbeans in Barbados, ASW= Americans of African ancestry in South western USA, PLCO= African-Americans from Eastern USA.

To better understand this pattern of *between-populations* homogenization of the African gene pool in the Americas, and for the geographic regions represented in our dataset for which there are historical demography records of origin and destination of Africans(1), we compared: (i) proportions of West-Central Africa-, Western Africa- and South-East Africa-associated ancestry clusters (Fig. 1) with (ii) expected proportion of these ancestry clusters, estimated from the proportions of arrivals from different locations (Fig. 4, *SI Appendix*, Table S5, S6 and section 5). Overall, for New World populations the proportions of South-Eastern African and Western African ancestry clusters are highly correlated with the expected ancestry based on the numbers of arrivals to Americas ports and departures from African ports (Spearman rho= 0.89, p = 0.02). However, for the West-Central African ancestry cluster the correlation does not reach significance (Fig. 4). For the entire American continent, we observe an excess of the observed West-Central Africa ancestry clusters (47.7% observed vs. 40% expected, being this a conservative estimation of the difference, see *SI Appendix*, section 5, p = < 2.2e-16), mainly determined by Southeastern Brazil and the US populations. The poorest concordance between observed and expected ancestries is observed in Southeastern Brazil, that presents more of the West-Central African ancestry cluster (37%) than expected (20%) (p = < 2.2e-16) and complementarily, less of the South/East African ancestry cluster than expected based on arrivals (55% observed vs. 76% expected (p < 2.2 × 10^−16^). The US population also shows an excess of the West-Central African ancestry cluster (54.7% observed vs. 43.1% expected, p < 3.33^−16^), compensated by a deficit of the Western African ancestry cluster (19.1% observed vs. 27.7% expected, p< 3.33^−16^). Therefore, the *between-population* homogenization of the African gene pool in the Americas is partly explained by the excess of the West-Central Africa ancestry cluster in Southeast Brazil and in the US (Fig. 4). The higher between-population homogeneity of the African gene pool in the Americas contributes to a more statistical power of genetic association studies involving individuals with African ancestry from different populations of the Americas.

**Fig. 4.**
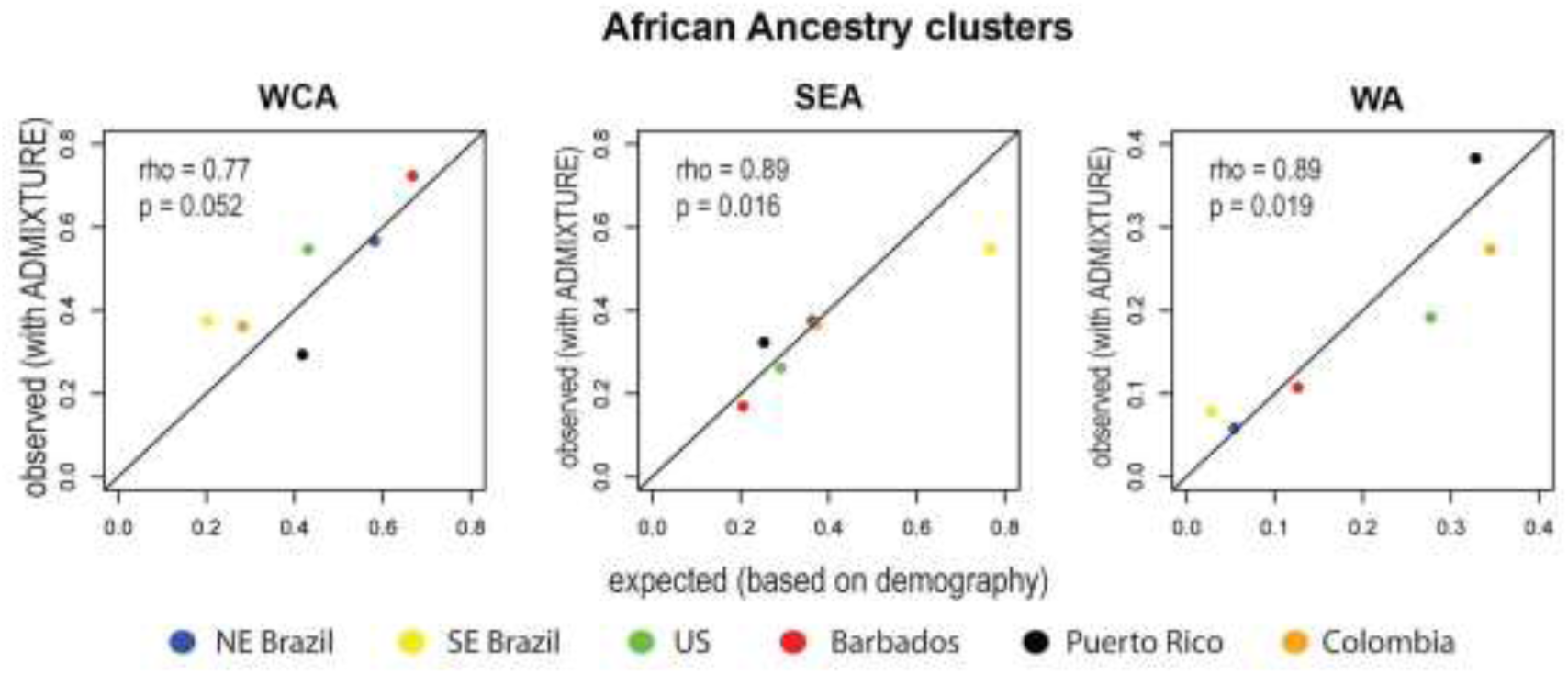
Observed and expected proportions of genomic African ancestry clusters in the Americas. We compared (i) the observed proportions of genomic African ancestry clusters (inferred using ADMIXTURE [Alexander et al. 2009]) in the vertical axis, with (ii) expected proportions of genomic African ancestry clusters, estimated based on demographic historical records from the African Voyages Database^1^, in the horizontal axis (see *SI Appendix* and Table S3 for details). rho: Spearman’s coefficients of correlation, p: significances. The significance was evaluated using randomization tests of 10,000 replications. WCA: West-Central Africa, SEA: South/East Africa, WA: Western Africa.

In conclusion, genetic data trace the African genetic roots of admixed individuals of the Americas to a broad geographic extension (from Western Africa to East Africa), associated with a high linguistic diversity (Niger Kordofanian non-Bantu and Western- and Eastern-Bantu language speakers). Considering the level of *between-populations* genetic differentiation in the African regions of origin of slaves, historical facts that homogenized the *between-populations* component of genetic diversity have predominated over facts that tend to maintain or increase it. This latter group of facts includes geographic (i.e. inter-continental distances and maritime winds/currents) and geopolitical factors (i.e. specific European colonial influences and possessions and the abolition of the slavery by British in 1807), that shaped an association of Western African ancestry with northern latitudes and South/East African ancestry with southern latitudes. Contrastingly, the following facts have contributed to gene flow between individuals with different African ancestries and thus, to the *between-populations* homogenization of the African gene pool in the Americas: (i) despite their specific European origins, traders/vessels transported slaves, frequently illegally, to different American continent ports (1, 18, 21); (ii) *forced amalgamation*, which is the preference of slave owners for slaves from different geographic and linguistic origins, so that they could not understand each other, and thus, reducing the risk of riots (22); and (iii) the role of islands such as Jamaica and Barbados, that centralized parts of arrivals of African slaves, and re-distributed them to different parts of the Americas (2). Moreover, the *between-population* homogenization of the African gene pool in the Americas is partly explained by the excess of the West-Central Africa ancestry cluster (the most prevalent in the Americas) in US and Southeast Brazil respect to demographic expectations, which suggest a spread of this ancestry in the American continent. Interestingly, in most of the Americas, the arrival of the largest contingent of Africans between 1,700 and 1,850 was almost synchronic with the intensification of admixture, which implies that this time interval was critical to shape the structure of the African gene pool in the New World. This study contributes with a population genetics perspective to the ongoing social, cultural and political debate regarding ancestry, race, and admixture in the Americas (23, 24).

## Methods

We analyzed a dataset including 6,267 unrelated individuals with more than 10% of African ancestry for 533,242 SNPs (*SI Appendix*, Table S1). We inferred population structure and admixture using ADMIXTURE (12) and Principal Component Analysis (25) for unlinked SNPs and ChromoPainter and fineSTRUCTURE (13) for haplotype-based analyses. Admixture dynamics was inferred using GLOBETROTTER (14). Demographic information of embarked and disembarked African slaves was obtained the African Voyages database (http://www.slavevoyages.org/voyage/search). Genetic differentiation between populations considering only the African gene pool was estimated using the *African-ancestry genetic distance* (AAGD, *SI Appendix*, section 4.1). Flowcharts and masterscripts of the analyses are available in the EPIGEN Scientific Workflow (26) web (http://ldgh.com.br/scientificworkflow). Details of Methods are in the SI Appendix.

Data Availability: EPIGEN-Brazil data are deposited in the European Nucleotide Archive (PRJEB9080 (ERP010139), accession no. EGAS00001001245, under EPIGEN Committee Controlled Access mode. The Nilotics and Kwa/Gur datasets are deposited in dbGaP at phs001705.v1.p1 and phs000838.v1.p1, respectively. The Botswana and Tanzania datasets from Sarah Tishkoff Lab are available at dbGaP accession number phs001396.v1.p1 and SRA BioProject PRJNA392485.

## Acknowledgments

We thank Sergio D Pena, Marcia Beltrame, Rosangela Loschi, Eduardo F Paiva, Fabrício Santos, Renan Souza, Claudio Struchiner, Ricardo Santos and Garrett Hellenthal for advice, discussions and criticisms.

## Author contributions

The project was conceived by MHG and ETS. MHG assembled datasets. MHG, VB, TPL, RGM, MMA, GSA, NMA, FSGK, MM, WCSM, LAM, MRR, HPSA, MLS, MOS, GSS, CZ analyzed genetic data. RL and RZ performed laboratory experiments. ETS supervised bioinformatic and statistical analyses. MD, RHG, HG, ACP, MFLC, MLB, BLH, SMM, SJC, SAT and MY contributed with data. MHG, VB, AWB, MY, SAT contributed to data interpretation. MHG, VB and ETS wrote the manuscript. All authors read the manuscripts and contributed with suggestions.

