## Supplementary_Materials for "Origins, admixture dynamics and homogenization of the African gene pool in the Americas"

**Summary**

**1. Methods** **3**

1.1. Populations 3

1.2. Data quality control 3

1.3. Relatedness 3

**2. Ancestry correspondence between African and admixed American continent populations, and the influence of geography and geopolitics.** **4**

2.1. Population structure analysis using genotype data 4

2.2. Population structure analysis using haplotype data 4

2.3. Chromosome painting 5

2.4. Haplotype based population substructure of the parental populations (African, European, Native American): fineSTRUCTURE 6

2.5. Ancestry contributions of African populations using haplotype-based methods 6

2.6. Supplementary Results 7

2.6.1. Continental and Subcontinental population structure 7

**3. Admixture dynamics in the Americas during the African diaspora** **8**

3.1.Supplementary results 10

**4. The genetic differentiation of the African gene pool in the Americas** **10**

4.1. African-ancestry genetic distance (AAGD) based on global ancestry inferences 10

**5. Comparison of ancestry observed results with expectations based on historical demography data of the African diaspora.** **11**

**6. References** **26**

### 1. Methods

#### 1.1. Populations

Our initial dataset included 10,155 individuals from North, Central and South America, Europeans, Native Americans and Africans from Western, Southern and Eastern Africa (Extended Data Table 1). The admixed American continent populations included: three Brazilian population-based cohorts from three different regions of Brazil (Salvador in Northeast, Bambuí in Southeast and Pelotas in South, from the EPIGEN-Brazil project [1] (https://epigen.grude.ufmg.br/); African Americans from the PLCO (Prostate, Lung, Colorectal and Ovarian Cancer screening) project [2] of the National Cancer Institute (NCI); Afro-Peruvians from the Instituto Nacional de Salud (INS, http://www.ins.gob.pe/) in Peru [3]; and the admixed American continent populations from the 1000 Genomes Project Phase III [4] [Americans of African Ancestry from US Southwest (ASW), African Caribbeans from Barbados (ACB), Colombians from Medellin, (CLM) and Puerto Ricans (PUR)].

As reference parental populations , we included 11 African populations from three sources: (i) all African populations from the 1000 Genomes project phase III [4] ; (ii) Botswana and Tanzania populations from the Tishkoff Laboratory [5]; and (iii) Ghanaian individuals from the Ghana Prostate Study Accra (from the NCI) and Ugandan individuals from the Epidemiology of Burkitt's Lymphoma in Eastern-African Children and Minors (EMBLEM) from the NCI [6].

#### 1.2. Data quality control

For Quality Control (QC) and merging of the genotype datasets, we followed the procedures described in Kehdy et al. [1] and in the workflows and MasterScripts ("Cleaning Process", "Quality Control - SQDC" and "Merging genotype datasets") of the EPIGEN-Brazil Scientific Workflow [7] (http://ldgh.com.br/scientificworkflow).

#### 1.3. Relatedness

Kinship coefficients between individuals ($\Phi ij$) of our admixed American continent populations were estimated using the method implemented in the Relatedness Estimation in Admixed Populations (REAP) software [8]. The estimates of $\Phi ij$ by REAP consider the individual ancestry proportion (IAP) from K parental populations and the K parental population's allele frequencies per SNP (KAF). We calculated the IAP and KAF using the ADMIXTURE software [9] in unsupervised mode assuming three clusters of ancestral populations (K = 3, Europeans, Africans and Native Americans). To estimate $\Phi ij$ among individuals of the parental populations, we used PLINK identity-by-descent estimation following the PLINK LD-based pruning recommendations [10].

To reduce the level of family structure within our study population samples while avoiding the loss of large number of related individuals, we used our method based on complex networks [1,11]. It represents families as groups of individuals (vertices) linked by $\Phi ij$ higher than an established cut-off (edges). When all families have this representation, they form a network of connected individuals. Here, we used the $\Phi ij$ cut-off $\geq 0.06$ to define family connections, in which all third, second and first-degree relatives are linked creating family clusters in the network. Then, we interactively excluded the most central individuals (i.e. with greatest number of relationship links) in the network, dissolving the family structure until all relationship links $\Phi ij \geq 0.06$ are disconnected. This process was repeated for each population (admixed and parental) separately to create a dataset comprised of 9,404 "unrelated" individuals.

### 2. Ancestry correspondence between African and admixed American continent populations, and the influence of geography and geopolitics.

Considering our main goal is to infer African ancestry, we restricted our analyses to individuals with more than 10% of African ancestry (estimated from ADMIXTURE results [K=3]). Then, we merged the remaining "unrelated" individuals from each population creating a Dataset U, comprised of 6,267 unrelated individuals for 533,242 shared SNPs, which was used in the following analyses.

#### 2.1. Population structure analysis using genotype data

For the population structure analyses, we used the workflows and MasterScripts ("Ancestry" and "Smart Eigenstrat") of the EPIGEN-Brazil Scientific Workflow [7] (http://ldgh.com.br/scientificworkflow). First, from the Dataset U we created the working Dataset GU (Genotype data of 208,400 SNPs and 6,267 Unrelated individuals) by pruning the SNPs in high linkage disequilibrium (LD) ($r2 \geq 0.4$), considering that ADMIXTURE and PCA assume that individuals are unrelated and linkage equilibrium among the markers [9,12]. Then, we used the dataset GU for unsupervised ADMIXTURE analysis (from K=2 to K=10, calculating cross-validation errors, Extended Data Figure 1) and for Principal Components Analysis (PCA - Extended Data Figure 2) [12,13]

#### 2.2. Population structure analysis using haplotype data

We applied haplotype-based methods (CHROMOPAINTER/fineSTRUCTURE/ GLOBETROTTER analyses [14,15]) to infer which African populations contributed to the populations in the Americas.

First, we phased the dataset U using the SHAPEIT software [16] using default parameters: 100 conditioning states and 20 main iterations of MCMC (Master script "Phase Data" from the EPIGEN-Brazil Scientific Workflow, http://ldgh.com.br/scientificworkflow). For haplotype-based methods, we did not exclude SNPs in linkage disequilibrium (LD) because the LD increases the power of the methods to detect subtle population structure [14]. Considering that the haplotype-based methods are computationally demanding (computational resources increases quadratically with the sample size), and that the results could be influenced by differences in samples sizes [14], we randomly sampled up to 50 unrelated individuals from each African, European and Native American population when the sample size was >50, and up to 200 unrelated individuals from each admixed American continent population. All processes involving these methods were performed following the flowchart "Haplotype-based methods" from the EPIGEN-Brazil Scientific Workflow, http://ldgh.com.br/scientificworkflow).

In summary, we created two datasets: (i) For genotype-based analyses using the ADMIXTURE method [9] and Principal Component Analysis [12], we used a dataset of 6,267 unrelated individuals analyzed for 208,400 unlinked SNPs ($r2 < 0.4$) shared by all 22 populations; (ii) for haplotype-based analyses with fineSTRUCTURE and GLOBETROTTER methods [14,15], we used original dataset which includes 533,242 SNPs shared by all the 22 populations and genotyped in the 6,267 individuals, but for some analyses we used a subset of individuals.

### 2.3. Chromosome painting

The haplotype based analyses have it basis on the generation of interindividual coancestry matrices of DNA chunks shared among individuals, generated by a process called *chromosome painting*, implemented in the CHROMOPAINTER software [14]. CHROMOPAINTER detects patterns of shared haplotypes between individuals by reconstructing individual haploid genome of putative recipients, as DNA chunks shared with other individual haploid genomes (the putative donors). In this process, each individual may be defined as recipient or donor (or both). CHROMOPAINTER uses a Hidden Markov Model to reconstruct each individual genome given: (i) the genomic information from other donor individual's phased data, and (ii) estimates of the two scaling parameters: recombination parameter and the mutation parameter, which depend on the studied populations, and are estimated by a previous run of CHROMOPAINTER on a subset of the dataset.

The *chromosome painting* produces two matrices of coancestry between individuals: (i) one based on the number of shared DNA chunks (chunkcounts) and (ii) one based on the sum of lengths of shared DNA chunks (chunklengths). Importantly, the coancestry matrices are dependent on the definition of individuals as recipients or donors, which depends on the hypothesis of gene flow to be tested.

Hereafter we describe how we use differently defined coancestry matrices to: assess the population structure of parental populations using the fineSTRUCTURE method [14] (Section 2.4); quantify the ancestry contributions of the parental populations to the admixed populations of the Americas using the MIXTURE MODEL regression implemented in GLOBETROTTER software [17,18] (Section 2.5); and to infer the admixture dynamic using also GLOBETROTTER (Section 3).

### 2.4. Haplotype based population substructure of the parental populations (African, European, Native American): fineSTRUCTURE

We assessed the population structure of parental populations by assessing if individuals that form genetically homogeneous clusters match their populations of origin. We restricted the analysis to African, European and Native American individuals and run CHROMOPAINTER allowing all them as both donors and recipients (-a flag of the software command).

To infer the scaling parameters (recombination scaling and mutation parameters), a preliminary CHROMOPAINTER run was performed with the same configuration of donors and recipients to be used for the fineSTRUCTURE analysis. For this purpose, we used 10 iterations of expectation–maximization, a subsample of the dataset (15 individuals per each population population) and a subsample of chromosomes (1, 7, 15 and 22). The resulting values for the parental populations were 267 for the 'recombination scaling constant' and 0.00043 for the mutation parameter.

We used the chunkcounts matrix resulting from the CHROMOPAINTER analysis to perform the model-based Bayesian clustering implemented in fineSTRUCTURE [14]. We ran fineSTRUCTURE in two steps: (i) to identify the clusters, we ran fineSTRUCTURE for 20 million iterations; the first ten million were burn-in iterations and the other ten million were used to sample the cluster configuration identified every 10,000 iterations. (ii) To identify the relationships among the clusters in a tree representation, we ran fineSTRUCTURE using 100,000 hill-climbing moves. This second step uses the clusters identified in the MCMC iteration with the highest posterior likelihood (step i) and seeks for a better configuration of the clusters. Then, we used the inferred fineSTRUCTURE tree to identify the homogeneous parental clusters of individuals. Because there is a very high correspondence between fineSTRUCTURE clusters and sampled populations (see results below), the former may be used as surrogate of the latter for the GLOBETROTTER inferences (Extended Data Figure 3).

### 2.5. Ancestry contributions of African populations using haplotype-based methods

We estimated the ancestry contribution of each African, European or Native American to the admixed American continent populations, using the MIXTURE MODEL regression implemented on GLOBETROTTER [15] (Figure 2B). For this purpose, the parental populations were set as donors, considering the high correspondence between the fineSTRUCTURE clusters and the corresponding populations. However, we excluded three Native Americans individuals that showed some level of European admixture and do not clusterized with their corresponding populations.

For the MIXTURE MODEL estimates, we used a different configuration of donors/recipients individuals for the *chromosome painting* setting process respect to that defined for the fine-STRUCTURE analysis. For the CHROMOPAINTER run for MIXTURE MODEL, we set individuals from the admixed American continent populations as recipient and individuals from the parental populations as both donors and recipients, and we applied the same scalar parameter values used for the *chromosome painting* of the parental individuals: 267 for the 'recombination scaling constant' and 0.00043 for the mutation parameter.

The MIXTURE MODEL uses the interindividual chunklengths co-ancestry matrix output from CRHOMOPAINTER. The chunklenghts co-ancestry matrix is constructed from *copyvectors* of admixed individuals (recipients), in which each element of a vector is the genome-wide length of all haplotypes shared with a specific donor. In order to infer the genetic contribution of the parentals populations (i.e. in this case the fineSTRUCTURE clusters) into each admixed population, the MIXTURE MODEL estimates the genome-wide average proportion that individuals share with each parental population. Then, using a non-negative least square regression, the MIXTURE MODEL estimates the regression coefficients by setting the average *copyvector* of the admixed population as an outcome variable and the *copyvectors* of the donors as predictor variables. Furthermore, the inferred coefficients are normalized to sum to unity. These coefficients are interpreted as the average contribution of a specific parental population to an admixed population. The inferred source populations were described with the suffix "-like", to emphasize that these populations are present day surrogates of the past real sources.

### 2.6. Supplementary Results

#### 2.6.1. Continental and Subcontinental population structure

The ADMIXTURE results are presented in Figure 1 and Extended Data Figure 1 and Extended Data Table 1, where K=6 corresponds to one of the lowest cross-validation errors. Furthermore, the Principal component (PC) analysis (Extended Data Figure 2 and Extended Data Table 1) showed similar pattern to ADMIXTURE.

The global pattern of ancestry inferred by ADMIXTURE K=6 [9] identified, in addition to the European and Native American continental clusters, four African genomic ancestry clusters (Fig. 1, Extended Data Table 1): West Central Africa, West Africa, Southern/Eastern African and Northern Ugandan, named on the basis of their association with African regions and populations. The Northern Ugandan ancestry cluster (cyan in Fig. 1) is highly associated with Nilotic African populations from Northern Uganda [6], but accounts for a very low proportion of the African ancestry in the Americas. In contrast, the other three African ancestry clusters were observed at high prevalence both in Africa and the Americas. Specifically:

(i) West-Central African (WCA, blue in Fig. 1), associated with Kwa and Gur ethnolinguistic groups in Ghana and with populations speaking Yoruba and Esan languages in Nigeria. These populations speak non-Bantu languages from the Niger-Kordofanian linguistic family (Extended Data Table 1). Notably, the Barbados African Caribbean population comparatively shows the highest contribution of Ghana-like populations (Kwa/Gur-like, 38%) as compared to other admixed populations of the Americas (Fig. 2, Extended Data Table 3).

(ii) Western African (WA, purple in Fig. 1), associated with the Mandinka population from Gambia and the Mende population from Sierra Leone. These populations speak non-Bantu languages from the Niger-Kordofanian Mande linguistic sub-family.

(iii) Southern/Eastern African (SEA, yellow in Fig. 1), associated with Herero and Mbukushu populations from Southern Africa, that speak Western Bantu languages, as well as with the Tswana population from Botswana (Southern Africa) and, with minor strength, with the Luhya population from Kenya. Tswana and Luhya speak Eastern Bantu languages. Bantu languages also belong to the Niger-Kordofanian linguistic family.

ADMIXTURE reported a differentiation between these four African genomic ancestry clusters that correspond to $F_{ST}$ values ranging between 0.01 and 0.025 (Extended Data Table 2).

We refined the ADMIXTURE results (based on allele frequencies) by performing haplotype-based analyses implemented in CHROMOPAINTER/fineSTRUCTURE/GLOBETROTTER [14,15] using the dataset U (Figure 2A and B). By performing fineSTRUCTURE analysis, our tree (Fig. 2A and Fig. 3) identified 16 genetically homogeneous clusters for a total of 609 individuals with higher correspondence between the clusters and populations, with some exceptions: 13 individuals of 609, clustered with individuals from other closely related populations. These individuals were Native Americans with evidence of European admixture (2 Ashaninkas and 1 Aymara), Tswana (2 out of 14 individuals), Herero (5 out of 37 individuals), Uganda (1 out of 49 individuals), Yoruba (1 out of 50 individuals), and Esan (1 out of 50 individuals). For the following analyses, we only include all parental individuals except for three Native Americans that showed European admixture.

Results from fineSTRUCTURE haplotype-based analysis and ADMIXTURE are consistent (Figs. 1, 2A; Extended Data Figure 3; Section 2): clusters defined by the latter method recapitulate the correlation of WCA, WA or SEA ADMIXTURE clusters with geography.

### 3. Admixture dynamics in the Americas during the African diaspora

For the admixture dynamics inference we used the GLOBETROTTER method [15], that uses the pattern of linkage disequilibrium among loci generated by admixture, as well as the dynamics of its decay, to infer: (i) the ancestry composition of the putative parental populations and (ii) the date the admixture event. This inference is performed by evaluating the fit of the observed data to five models that represent five admixture scenarios: no admixture, uncertain, one date, one date multiway and two dates. Operationally, the GLOBETROTTER method [15] requires the information of the chunklengths coancestry matrix (copyvectors) and inferences of the donor haplotype for each recipient locus (painting samples). With this information, GLOBETROTTER uses the relationship between the probability of finding two chunks of different donors as a function of the genetic distance along the genome of individuals from a target admixed population. This relationship for each pair of putative parental populations, represented as *coancestry curves*, provides information about when the admixture events occurred and about the genetic composition of the parental populations.

GLOBETROTTER requires copyvectors and painting samples of the individuals of the admixed populations. The copyvectors information is the same used for the MIXTURE MODEL (Section

2.5). The painting samples are the inference of the possible donor (from parental populations) for each locus of recipient haplotype.

This inference is performed for each population and requires population specific recombination and mutation parameters. We calculated the scalar parameters for each recipient population of the Americas using 30 iterations of expectation-maximization for a subset of individuals and chromosomes (1, 7, 15 and 22) for each target population. Then, we performed CHROMOPAINTER analysis for each target population with the population-specific parameters, setting the target population as recipient and the parental clusters as donors. With the copyvectors and painting samples information, we did GLOBETROTTER analysis.

First, GLOBETROTTER tests the "no admixture model" and determine its p-value. If this model is rejected, GLOBETROTTER infers the parameters of the most plausible model based on the coancestry curves and threshold predetermined values. For this purpose we used the standardization implemented in GLOBETROTTER with the flags null.ind: 1. This standardization identifies the component of the co-ancestry curves that do not correspond to admixture LD, being more conservative [15] than the alternative flags null.ind: 0.To obtain confidence intervals of the inferred date of admixture we performed 100 bootstrap replicates (Fig. 2C and Extended Data Table 4).

GLOBETROTTER inferred dates are in number of generations (G). To calculate the admixture date in years we used the following formula : Y = A - G*L, where A is the mean year of birth of the individuals in the sample, G is the number of generations inferred by GLOBETROTTER and L is the length of a generation in years. We used L = 30 years. The A values were: 1929 for PLCO cohorts, 1980 for 1000 Genomes samples (ASW, ACB, PUR and CLM), 1980 for Afro Peruvians, 1998 for the Salvador cohorts, 1928 for the Bambui cohorts and 1982 for the Pelotas cohorts (Fig. 2C and Extended Data Table 4). For the plotting of GLOBETROTTER results, we used and modified the codes used by Busby et al. [19] which are available in https://github.com/georgebusby/admixture_in_africa. The modified version of the codes are available in as part of the EPIGEN Scientific Workflow (http://ldgh.com.br/scientificworkflow).

Furthermore, we compared demographic history and admixture dynamics considering the information of (i) the number of disembarked slaves by year from the African Voyages database (http://www.slavevoyages.org/voyage/search) and the (ii) dates of admixture inferred by 100 bootstrap replicates performed during GLOBETROTTER. We plotted these comparisons for six geographical regions with available data: United States, Barbados, Puerto Rico, Colombia, Salvador and Southeast Brazil (Extended Data Figure 4).

**3.1.Supplementary results**

We inferred the most likely Post-Columbian admixture events (dates and admixture sources, Fig. 2C and Extended Data Figure 4 and Extended Data Table 4) in the admixed American continent populations. The genetically homogeneous clusters of individuals (parental or donors) inferred by CHROMOPAINTER/fineSTRUCTURE analyses (Fig. 2A) were used as surrogates of populations. We only excluded 3 Native American individuals that clustered together and showed European admixture (Extended Data Figure 1 and 3). For our admixed American continent populations, the best fit of their co-ancestry curves were with the one-date and two-dates models (Extended Data Table 4).

In addition to the continental trend of admixture observed for the 1750-1850 interval, we noted that while the intensification of admixture was synchronic, the parental population contributions were different among the admixed American continent populations (Fig. 2C and Extended Data Table 4): (i) For Brazilians, Colombians and Puerto Ricans, the predominant European source was Iberic, while for former UK colonies it was of North European origin. Consistent with Harris et al. [3], Peru is an exception because it shows admixture between two already admixed populations, one with predominant Native American ancestry. Thus, the Afro-Peruvians are one of the only two studied Afro-Native American populations of the American continent, together with the Honduras Garifuna[20]. (ii) Former Spanish colonies (Puerto Rico, Colombia and Peru) and Southeastern Brazil populations show signatures of earlier or later intensification of admixture with respect to the 1750-1850 interval: in Southern Brazilian populations admixture intensified in the 100 years previous to the 1750-1850 interval. In the former Spanish colonies, admixture may have occurred as early as in the 16th Century, with Puerto Rico showing the earliest signal, which is interesting considering that the first slaves transported in the Americas arrived to the Caribbean region [21] .

### 4. The genetic differentiation of the African gene pool in the Americas

To estimate the genetic differentiation between the admixed American continent populations and between African populations, considering only the African gene pool, we used the following approach: We defined a genetic distance hereafter called *African-ancestry genetic distance* (AAGD) that, based only on the proportion of African ancestry clusters, solely refers to the African gene pool of the populations of the Americas (i.e. excluding the information derived from European and Native American ancestries (Section 4.1.).

#### 4.1. African-ancestry genetic distance (AAGD) based on global ancestry inferences

AAGD is based on:
(i) the mean proportions of the subcontinental African ancestry clusters from each population based on ADMIXTURE results (K=6) (Extended Data Table 1). In the case of population of the Americas, respect to the total African ancestry.
(ii) the $F_{ST}$ between the African ancestry clusters estimated by the ADMIXTURE software [9] (Extended Data Table 2).

AAGD between two populations is given by the sum of the Euclidean genetic distances between each pair of subcontinental ancestries weighted by the $F_{ST}$ (in sensu ADMIXTURE [9]) between the ancestry clusters.

Considering two populations (A and B) with three ancestry clusters (x, y and z), the African-ancestry genetic distance is calculated as follow

$$\text{Distance}\,(A_x,\ A_y,\ A_z,\ B_x\ ,\ B_y,\ B_z\,) = Fst_{x,y}\ \sqrt{(A_x - B_x)^2 + (A_y - B_y)^2}\ + Fst_{x,z}\ \sqrt{(A_x - B_x)^2 + (A_z - B_z)^2}\ + Fst_{y,z}\ \sqrt{(A_y - B_y)^2 + (A_z - B_z)^2}$$

Where $A_i$ is the population ancestry proportion of the ADMIXTURE cluster i, respect to the total African ancestry.

We compared the AAGD with adaptations of three classical population genetics measures: (1) Compute pairwise $F_{ST}$ [22]; (2) Slatkin's genetic distance [23] and (3) Reynolds's genetic distance [24] estimated using the Arlequin software [25]. To calculate these versions of classical genetic distances, we used the proportions of African ancestry clusters as being allele frequencies of a single locus. The correlation between the AAGD and the other three matrices (from the classical statistics) were > 0.76, p = 0.02, using the Mantel correlation test [26], implemented as in the "ade4" library for the R software (Extended Data Figure 5).

**5. Comparison of ancestry observed results with expectations based on historical demography data of the African diaspora.**

For the geographic regions represented in our dataset for which there are historical demography records of origin and destination of Africans (the Transatlantic Slave Trade database) [27], we compared: (i) the inferred WCA, WA and SEA genomic ancestries (Fig. 1 and Extended Data Table 1) with (ii) the expected proportion of these genomic ancestries, estimated from the proportions of arrivals from different locations (Fig. 3, Extended Data Table 5-6). We assumed that the genomic ancestry in current African populations are reasonable proxies of the populations during the Transatlantic Slave Trade and that, in general, as in most human populations, there is spatial autocorrelation of genetic structure in Africa [28].

We download the table of historical records from the African Voyages database (http://www.slavevoyages.org/voyage/search) using the criteria of search: "Embarkation Regions" in the columns, "Disembarkation Ports" in the rows and "Sum of embarked slaves" in the cells. For these comparisons, we used population proxies of destiny of the African diaspora (Extended Data Table 5). To represent these destiny proxies, we used our genomic data of current admixed American continent populations close to ports of disembark that are historically related to the African diaspora. As observed ancestry estimates, we used the proportion of ancestry estimated by ADMIXTURE (K = 6, Fig. 1 and Extended Data Table 1) of specific populations in the Americas as described in the Extended Data Table 5-6.

For the expected ancestry in a specific destiny proxy, we considered the proportion of individuals from the embarkation major regions (representing the ancestry origin proxies,

(Extended Data Table 6) that arrived in specific ports of disembarkation in the Americas using the data available in the African Voyages database [27]. To avoid the unrealistic assumption that the individuals from embarkation major regions have a homogenous ancestry, we calculated the weighted expected ancestry. This was estimated by taking into account the proportion of WCA, WA, SEA genomic ancestry clusters estimated in selected current African populations from the embarkation major regions that contributed to the African genomic pool in the Americas (Extended Data Table 6 and Figs. 1A and 2B): (i) Kwa/Gur and Yoruba populations for the WCA proportion; (ii) Mandinka (GWD) and Mende (MSL) for the WA proportion and (iii) Mbukushu and Luhya (LWK) for the SEA proportion. An exception were the Brazilian populations, that we used only GWD population to obtain the WA proportion, since the Mende (MSL) did not genetically contribute to the Brazilian populations (Fig. 2A and B, Extended Data Table 3).

Thus, we calculated the weighted expected proportion of the i-ancestry cluster ($Wi$) for each proxi-destiny population as $Wi = \sum_{p=1}^{n} e_p * o_{p,i}$,

where

***n*** is the number of African population proxies of origin (***p***);

$\boldsymbol{e_p}$ is the expected ancestry of the population proxies of origin ***p*** based on the proportion of individuals arrived from each of the n African populations, respect to the total of individuals disembarked in the population;

$\boldsymbol{o_{p,i}}$ is the observed proportion of the ADMIXTURE ancestry cluster ***i*** of population proxies of origin ***p***.

To assess the correlation between observed and expected ancestries, we applied the Spearman correlation test, implemented in the R and the significance was evaluated using 10,000 randomization tests using the following function:

```
randomize = function(a,b,nreps){
 r.obs = cor(a,b,method = "spearman")
 cat("The obtained correlation is ",r.obs,'\n')
 r.random = numeric(nreps)
 for (i in 1:nreps){
 Y = a
 X = sample(b, replace = FALSE)
 r.random[i] = cor(X,Y, method = "spearman")
 }
 prob = length(r.random[r.random >= r.obs])/nreps
 cat("P-Value =",prob)

 hist(r.random, breaks = 100, main = expression(paste("Distribution around ",rho, "= 0")),
 xlab = "r from randomized samples")
```

}

The *a* and *b* are numeric vectors of data values and *nreps* is the number of randomization tests.

To compare the observed and the expected distributions of the proportion of individual African ancestry cluster (relative to the total African ancestry), we followed this procedure considering that for the expected distribution we only have its mean $W_i$ : (1) we assessed the observed distributions of the proportion of individual African ancestry cluster and their moments (mean and variance were sufficient in all cases) and used the R package *fitdistrplus* to fit them to theoretical distributions. In our cases, Normal for WCA and SEAB in Southeast Brazil and WA in US, Beta for WCA in Total America and Weibull for WCA in US. (2) To create the expected distribution of the proportion of African ancestry clusters, we sampled from the adjusted-theoretical distribution with the expected mean and the same variance that the observed distribution, conditioning on the same number of individuals than the observed distribution. (3) We compared the observed and expected distributions by using the non-parametric Kolmogorov-Smirnov test (ks.test, R Base Package). Results of these comparisons are in Extended Data Table 7.

### Extended Data Figures

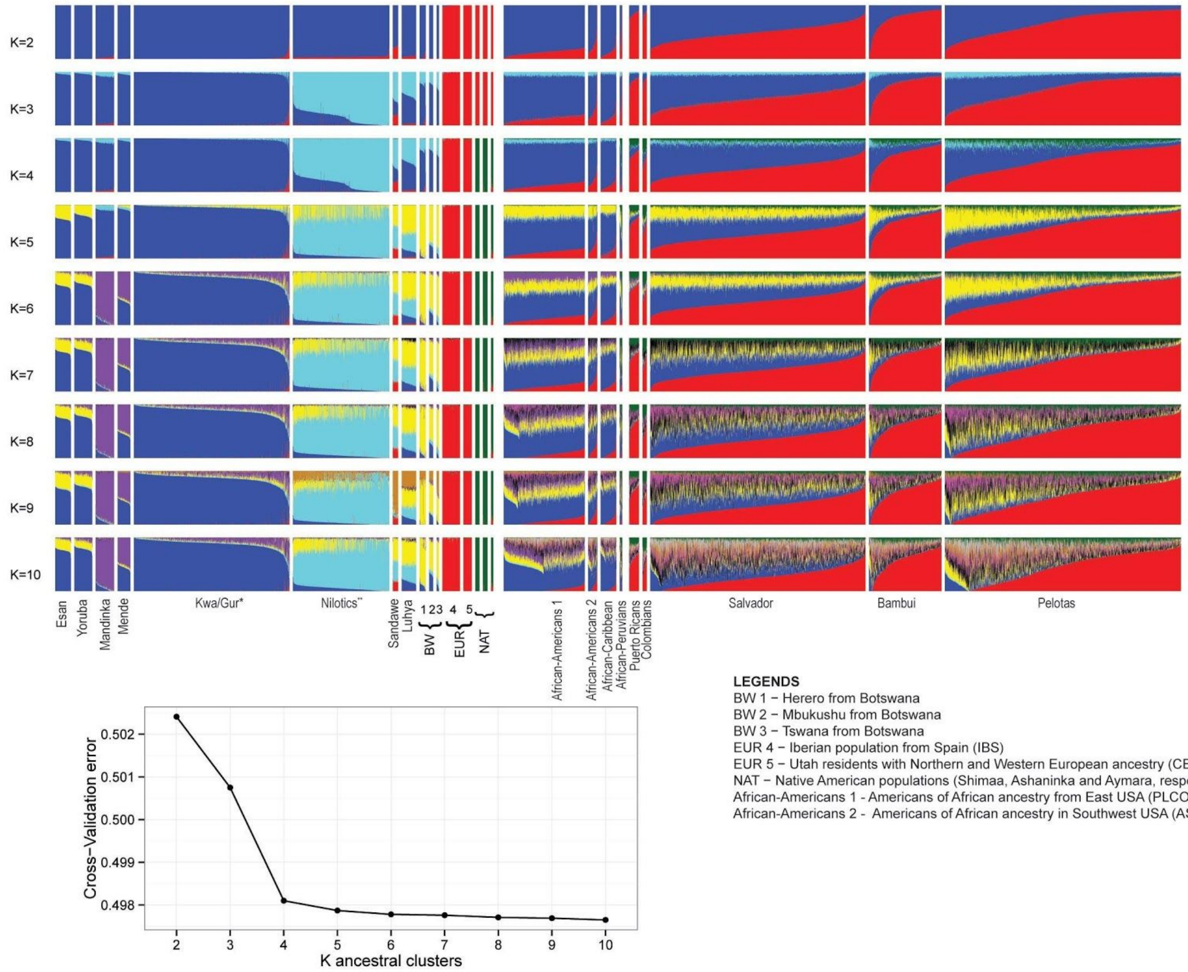

**Extended Data Figure 1. Barplot representation of the individual ancestry proportion of unrelated individuals inferred using ADMIXTURE.** The proportions of individual ancestry values were calculated using the number of parental K = 2 to K = 10 for the Dataset GU (Genotype dataset used to study the population structure). Ancestral populations are sorted so that each one is assigned to a specific group in this order: Africans, Europeans Native American, and admixed American continent populations. Each bar represents an individual and each color a specific ancestry cluster. Barplots are sorted for each K by decreasing amount of the blue ancestry cluster so that individuals are not vertically aligned across the Figure. We also represented the ADMIXTURE cross validation errors. *The Kwa/Gur dataset includes approximately 35 ethno-linguistic groups, predominantly from the Kwa and Gur Niger-Congo linguistic group[6]. **The Nilotics dataset includes predominantly three ethnolinguistic groups in Northern Uganda (Langi, Acholi and Lugbara) from the Nilotic linguistic group[6].

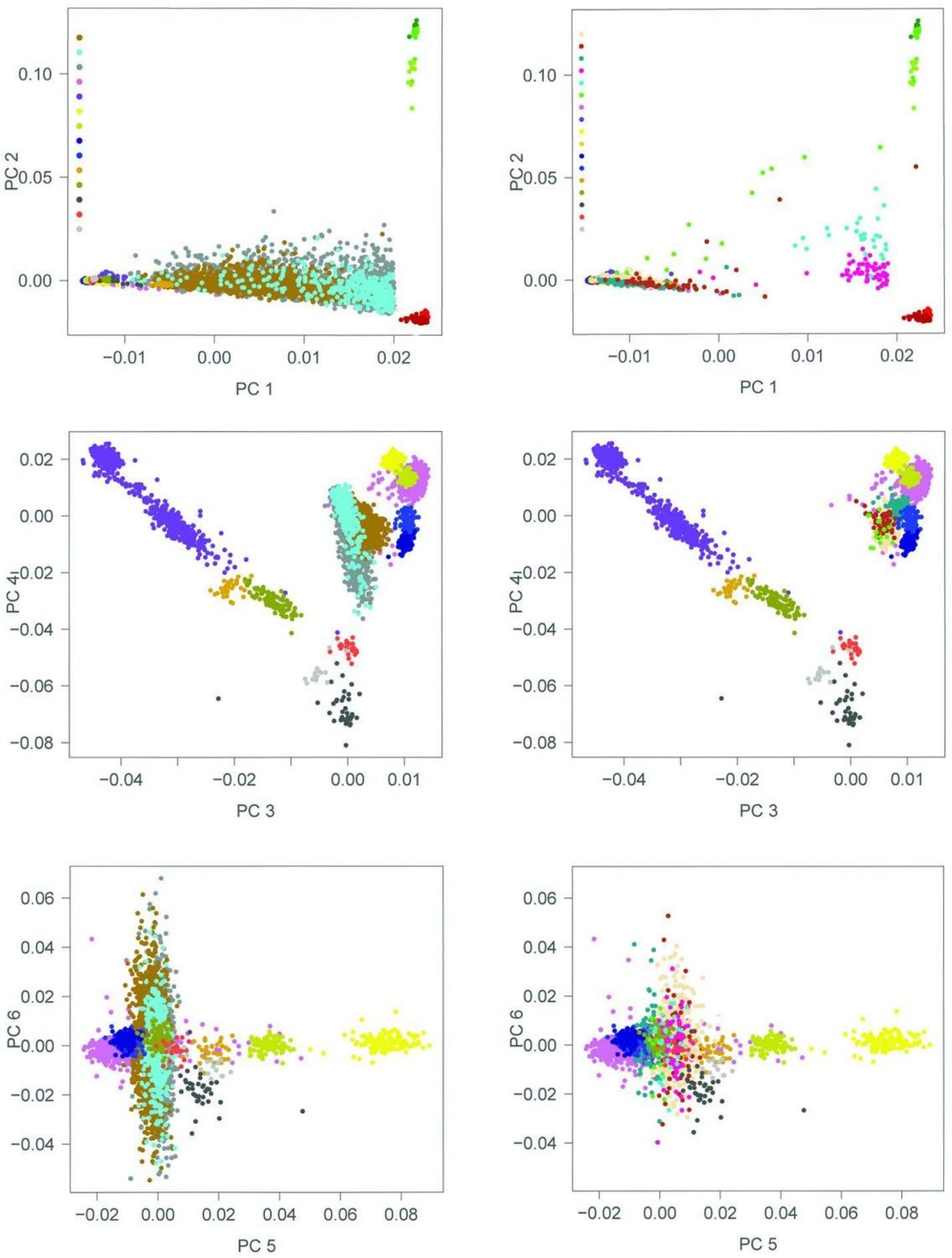

**Extended Data Figure 2. Principal Component Analysis (PCA).** PCA and the percentage of variability identified by each PC for Dataset GU (Genotype dataset used to study the population structure). For a better visualization of the individual's distribution along the PC axes, we represented the Brazilians with the parental populations in the left plot and the other admixed American continent population with the parental populations in the right plot. Black ellipses in the PC1 vs PC2 plot represent the Native Americans individuals (Ashaninka, Aymara and Shimaa) and European individuals (CEU=Utah Residents (CEPH) with Northern and Western Ancestry-USA and IBS=Iberian population in Spain). Plots of PC3 vs PC4 and PC5 vs PC6 reflect only African diversity, and therefore, we only plotted individuals from Africa and from the American continent. African-Americans 1 are the Americans of African ancestry from East USA (PLCO project). African-Americans 2 are the Americans of African ancestry in Southwest USA (ASW). *The Kwa/Gur dataset includes approximately 35 ethno-linguistic groups, predominantly from the Kwa and Gur Niger-Congo linguistic group[6]. **The Nilotics dataset includes predominantly three ethnolinguistic groups in Northern Uganda (Langi, Acholi and Lugbara) from the Nilotic linguistic group[6].

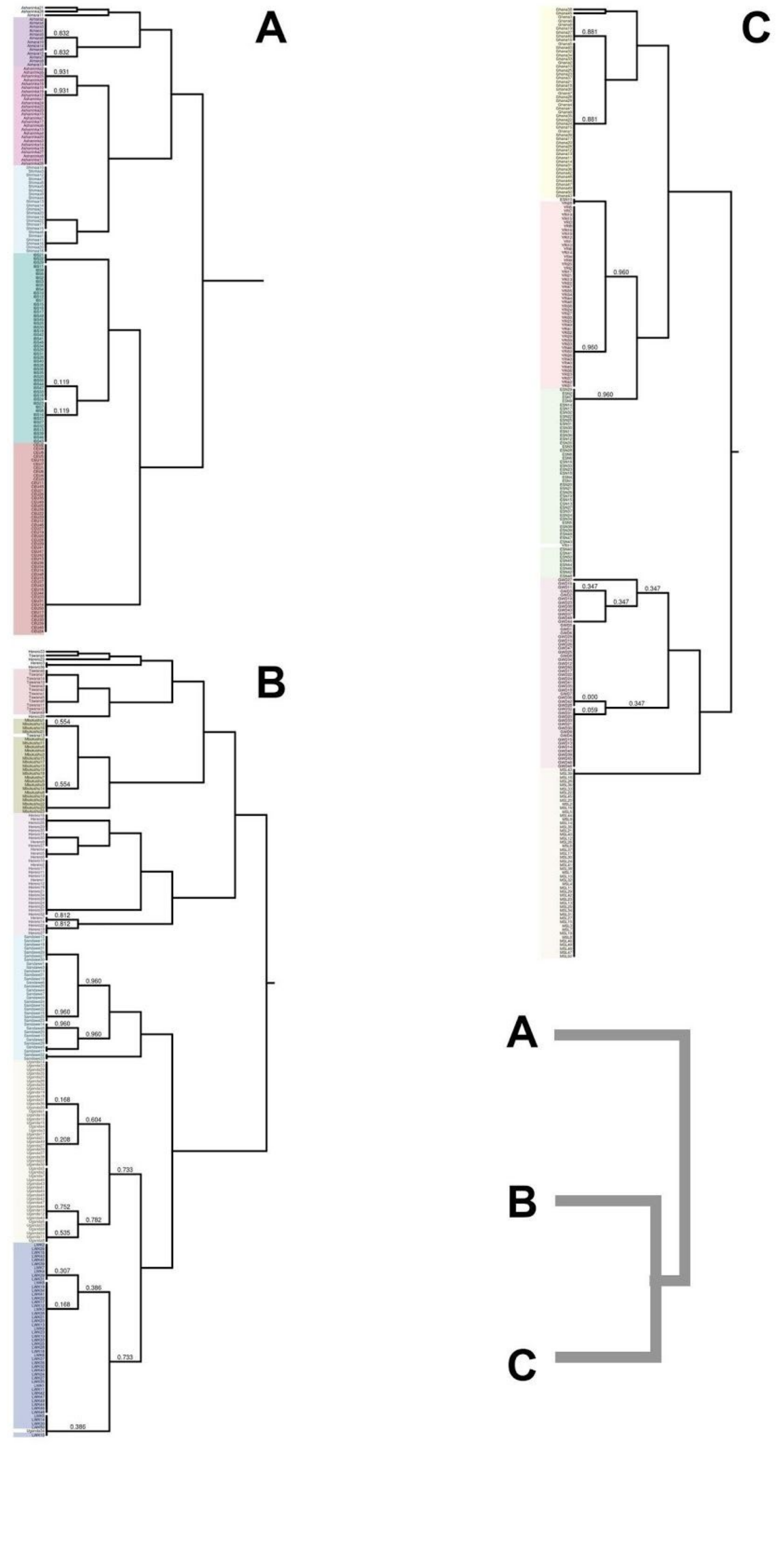
A
0.832
0.832
0.931
0.931
0.119
0.119
B
0.554
0.554
0.812
0.812
0.960
0.960
0.960
0.960
0.168
0.604
0.208
0.733
0.752
0.782
0.535
0.307
0.386
0.168
0.733
0.386
C
0.881
0.881
0.960
0.960
0.960
0.347
0.347
0.347
0.000
0.059
0.347
A
B
C

**Extended Data Figure 3. fineSTRUCTURE tree of the parental individuals.** Tree of the sampled individuals pooled in 16 worldwide clusters as inferred by fineSTRUCTURE. Inferred clusters are highly correlated with the studied populations. Shadow colors are related to each population. Individuals non included in shadow areas do not clustered with its nominal populations. (A) Cluster including Native Americans (Aymaras, Ashaninkas, Shimaa) and European (Iberian from Spain [IBS] and Utah residents with Northern and Western European ancestry [CEU]). (B) African cluster including Bantu from East Africa (Luhya from Kenya [LWK] and Sandawe) and South Africa (Herero, Mbukushu and Tswana), and Nilotics from Northern Uganda populations. (C) African cluster including West (Gambia [GWD] and Sierra Leona [MSL]) and West Central (Ghana [Kwa/Gur] and Nigeria[YRI and ESN]) populations**.**

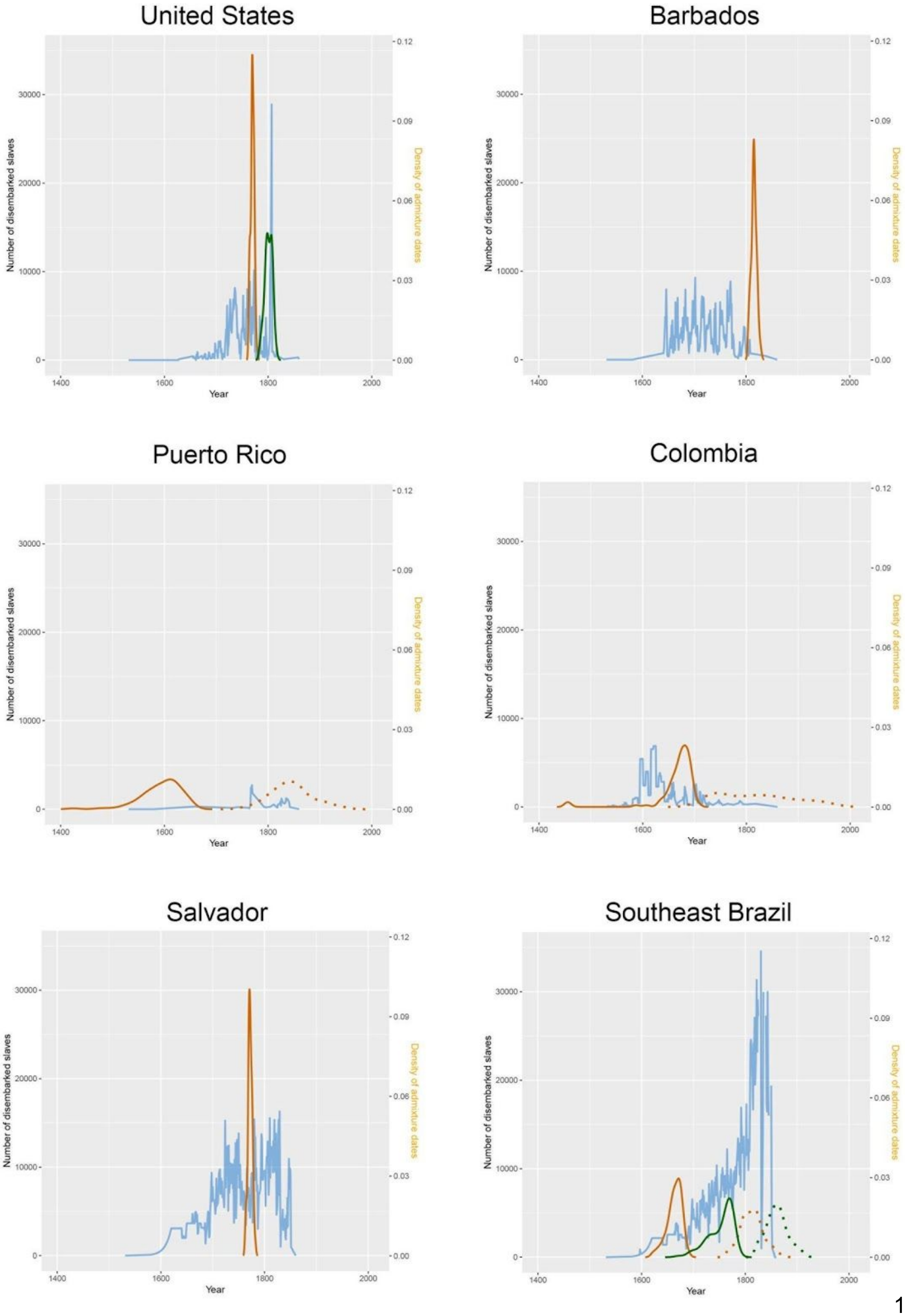
United States
Barbados
Puerto Rico
Colombia
Salvador
Southeast Brazil
Number of disembarked slaves
Density of admixture dates
Year

**Extended Data Figure 4. Relationship between number of Disembarked slaves and admixture dates inferred by GLOBETROTTER.** We plotted the distribution of the number of disembarked slaves (blue line) during the XV to the XVIII centuries in six ports along the Americas: United States, Barbados, Puerto Rico, Colombia, Salvador and Southeast Brazil. Overlapping this information, we plotted a smoothed distribution of the admixture dates inferred for 100 bootstrap replicates in GLOBETROTTER. For the United States, we plotted the admixture dates distribution for PLCO (green curve) and ASW (orange curve) populations. For the Southeast Brazil, we plotted the admixture dates distribution for Bambui (orange curve) and Pelotas (green curve) populations. For populations in which the two dates admixture model was inferred, the earlier and the recent date were plotted as unbroken and dashed lines, respectively.

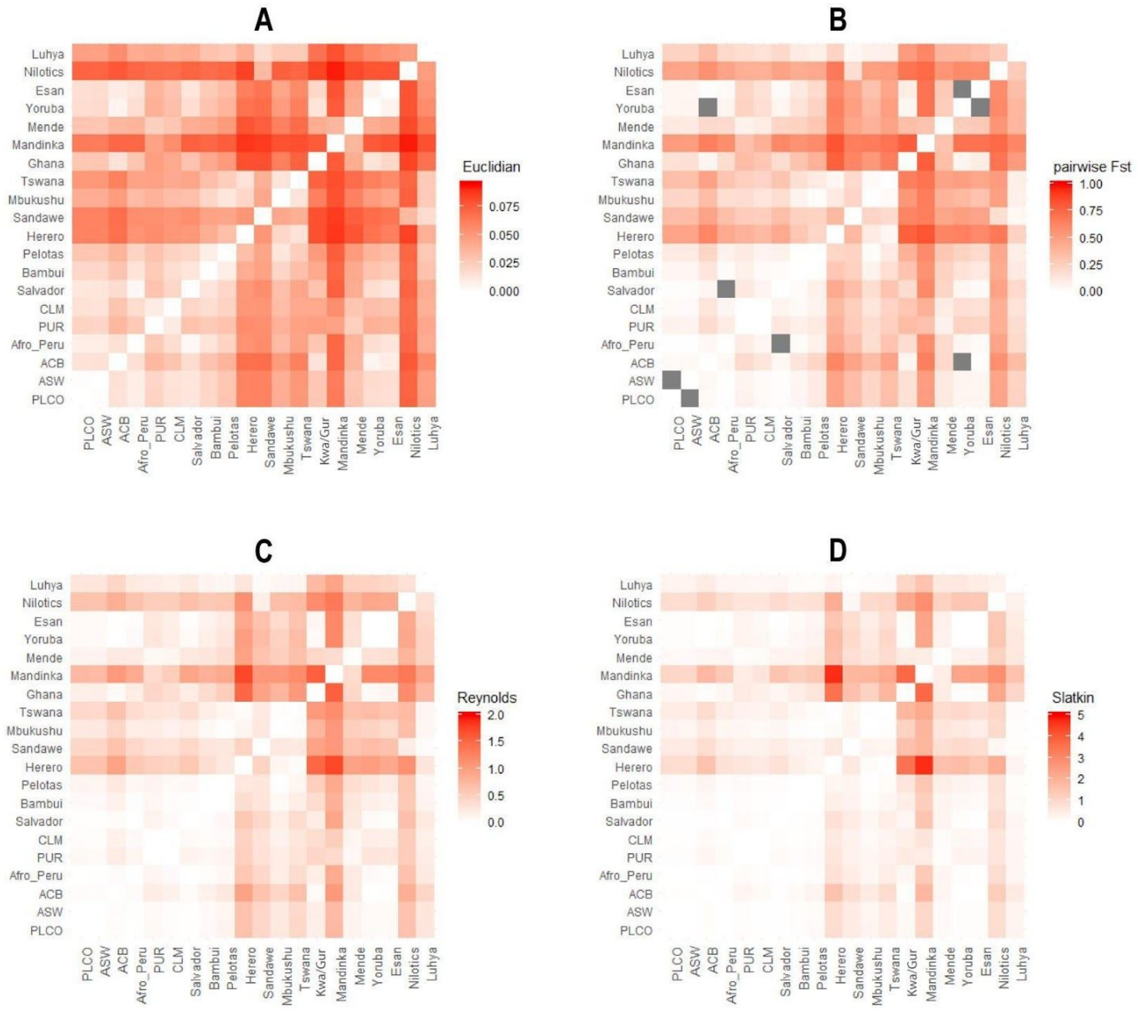

**Extended Data Figure 5. Pairwise genetic distances (based on global ancestry) of the African gene pool between populations of the admixed American continent and Africa.** Scatterplot Matrices of the: (A) Euclidean genetic distance (see SI text); (B) Compute pairwise differences (Nei and Li, 1979); (C ) Slatkin's genetic distance (Slatkin, 1995) and (D) Reynolds's genetic distance (Reynolds et al. 1983). All matrices showed a mantel correlation > 0.76. CLM=Colombians from Medellin, PUR=Puerto Ricans from Puerto Rico, ACB=African Caribbeans in Barbados, ASW= Americans of African ancestry in South west USA, PLCO= African-Americans from East USA.

**Extended Data Table 1. Populations included in this study and proportions of subcontinental African ADMIXTURE clusters, respect to the total African ancestry, in the admixed American continent and African populations.** Populations: CLM=Colombians from Medellin, PUR=Puerto Ricans from Puerto Rico, ACB=African Caribbeans in Barbados, ASW= Americans of African ancestry in South west USA,PLCO= Americans with African ancestry from East USA, CEU=Utah Residents (CEPH) with Northern and Western Ancestry-USA, IBS=Iberian population in Spain, YRI = Yoruba in Ibadan-Nigeria, ESN = Esan in Nigeria, GWD= Gambian in Western Divisions in the Gambia , LWK= Luhya in Webuye, Kenya, MSL= Mende in Sierra Leone, BWHE = Herero in Botswana, BWMB= Mbukushu in Botswana, BWTS = Tswana in Botswana, TZSW= Sandawe in Tanzania. African Ancestry cluster: WCA=West Central Africa; WA= West Africa; SEA= South-East Africa; NU= Nilotic Ugandan.

| Population | Country | Individuals (n) | Array/ Sequencing | SNPs | Subcontinental African ADMIXTURE clusters | | | | Ref * |
|---|---|---|---|---|---|---|---|---|---|
| | | | | | WCA | SEA | WA | NU | |
| **AMERICAN CONTINENT ADMIXED POPULATIONS** | | | | | | | | | |
| Salvador | Brazil | 1,309 | 2.5M | 2,306,356 | 0.567 | 0.338 | 0.058 | 0.037 | 1 |
| Bambui | Brazil | 1,442 | 2.5M | 2,306,356 | 0.407 | 0.442 | 0.089 | 0.062 | 1 |
| Pelotas | Brazil | 3,736 | 2.5M | 2,306,356 | 0.341 | 0.541 | 0.068 | 0.05 | 1 |
| Afro-Peruvian s | Peru | 15 | 2.5M | 2,306,356 | 0.557 | 0.309 | 0.093 | 0.041 | 2 |
| CLM | Colombia | 94 | WG | - | 0.362 | 0.312 | 0.273 | 0.053 | 3 |
| PUR | Puerto Rico | 104 | WG | - | 0.294 | 0.264 | 0.383 | 0.06 | 3 |
| ACB | Barbados | 96 | WG | - | 0.723 | 0.152 | 0.107 | 0.018 | 3 |
| ASW | U.S. | 61 | WG | - | 0.552 | 0.234 | 0.187 | 0.027 | 3 |
| PLCO | U.S. | 524 | MP | 1,022,207 | 0.542 | 0.232 | 0.196 | 0.03 | 4 |
| **EUROPEAN REFERENCE PARENTAL POPULATIONS** | | | | | | | | | |
| CEU | U.S. | 99 | WG | - | - | - | - | - | 3 |
| IBS | Spain | 107 | WG | - | - | - | - | - | 3 |
| **AFRICAN REFERENCE PARENTAL POPULATIONS** | | | | | | | | | |
| YRI | Nigeria | 108 | WG | - | 0.762 | 0.179 | 0.052 | 0.007 | 3 |
| ESN | Nigeria | 99 | WG | - | 0.726 | 0.25 | 0.022 | 0.002 | 3 |
| GWD | Gambia | 113 | WG | - | 0.06 | 0.007 | 0.923 | 0.01 | 3 |
| LWK | Kenya | 99 | WG | - | 0.14 | 0.517 | 0.013 | 0.33 | 3 |
| MSL | Sierra Leone | 85 | WG | - | 0.445 | 0.033 | 0.519 | 0.004 | 3 |
| Herero | Botswana | 76 | 5M | 4,192,458 | 0.051 | 0.896 | 0.024 | 0.03 | 5 |
| Mbukushu | Botswana | 44 | 5M | 4,192,458 | 0.228 | 0.668 | 0.039 | 0.064 | 5 |

| | | | | | | | | | |
|---|---|---|---|---|---|---|---|---|---|
| Kwa/Gur** | Ghana | 968 | 5M | 4,641,218 | 0.881 | 0.027 | 0.076 | 0.016 | 6 |
| Nilotics*** | Uganda | 808 | 5M | 4,641,218 | 0.045 | 0.15 | 0.013 | 0.792 | 7 |
| **NATIVE AMERICAN REFERENCE PARENTAL POPULATIONS** | | | | | | | | | |
| Shimaa | Peru | 45 | 2.5M | 2,348,597 | - | - | - | - | 8 |
| Ashaninkas | Peru | 44 | 2.5M | 2,348,597 | - | - | - | - | 8 |
| Aymaras | Peru | 16 | 2.5M | 2,348,597 | - | - | - | - | 8 |
| TOTAL | | 10,155 | | | | | | | |

*Datasets used in the study: 1 = Dataset from EPIGEN Consortium (Kehdy et al. 2015), 2 = Dataset from Instituto Nacional de Salud (INS) - Peru, 3 = Public dataset published by the 1000 genomes Project , 4 – PLCO (Prostate, Lung, Colorectal and Ovarian Cancer screening) project from National Cancer Institute (NCI-NIH) - U.S, 5 = Dataset from Sarah Tishkoff Lab - University of Pennsylvania (Crawford et al. 2017), 6 = Dataset from Ghana Prostate Study (NCI-NIH) and 7 = Dataset from EMBLEM (Epidemiology of Burkitt's Lymphoma in East-African Children and Minors - (NCI-NIH), 8 – Dataset from the Tarazona–Santos group LDGH (Laboratory of Human Genetic Diversity) dataset. **The Kwa/Gur dataset includes approximately 35 ethno-linguistic groups, predominantly from the Kwa and Gur Niger-Congo linguistic group[6]. ***The Nilotics dataset includes predominantly three ethno-linguistic groups in Northern Uganda (Langi, Acholi and Lugbara) from the Nilotic linguistic group[6].
The field array/sequencing has some abbreviations: 2.5M is an abbreviation for the Illumina Omni2.5, 5M is an abbreviation for the Illumina Omni5M, WG is an abbreviation for 1KGP, MP is an abbreviation for Multiple Datasets, which is composed of 1M/550K/610K/660W/Hap1/OmniX (illumina and affymetrix)

**Extended Data Table 2. Genetic differentiation (FST) matrix among ADMIXTURE ancestry clusters obtained with K=6.** The blue, yellow, purple, cyan, red and green colors refer to West-Central, South/East, Western, Northern Ugandan, European and Native American ancestry clusters.

| Ks | | | | | |
|---|---|---|---|---|---|
| | 0.018 | | | | |
| | 0.010 | 0.013 | | | |
| | 0.024 | 0.025 | 0.021 | | |
| | 0.118 | 0.120 | 0.119 | 0.115 | |
| | 0.208 | 0.209 | 0.208 | 0.205 | 0.155 |

**Extended Data Table 3. MIXTURE MODEL results for the ancestry contribution of Parental populations into admixed American continent populations and the relative contribution of the African ancestries.** We infer the contribution of the 11 African and three Native American and two European populations. Moreover, the column “African proportion” shows the total African ancestry in the genome of the admixed populations. Finally, column names with the prefix “RAP” for Relative African population is for the decomposition of the total African ancestry of admixed populations of the Americas in their sub-continental proportions.

| | Sandawe | Tswana | Herero | Mbukushu | LWK | Nilotics** | MSL | GWD | Kwa/Gur* | ESN | YRI | IBS | CEU | Ashaninkas | Aymaras | Shimaa |
|---|---|---|---|---|---|---|---|---|---|---|---|---|---|---|---|---|
| ACB | 0.001 | 0.003 | 0.003 | 0.015 | 0.030 | 0.002 | 0.059 | 0.026 | 0.336 | 0.063 | 0.345 | 0.000 | 0.113 | 0.000 | 0.002 | 0.000 |
| PUR | 0.004 | 0.002 | 0.003 | 0.017 | 0.020 | 0.003 | 0.012 | 0.054 | 0.021 | 0.014 | 0.041 | 0.635 | 0.036 | 0.038 | 0.090 | 0.012 |
| PLCO | 0.005 | 0.011 | 0.011 | 0.052 | 0.058 | 0.018 | 0.079 | 0.087 | 0.169 | 0.098 | 0.295 | 0.007 | 0.101 | 0.002 | 0.006 | 0.001 |
| ASW | 0.003 | 0.009 | 0.009 | 0.041 | 0.062 | 0.008 | 0.076 | 0.081 | 0.119 | 0.083 | 0.281 | 0.007 | 0.190 | 0.007 | 0.022 | 0.002 |
| CLM | 0.003 | 0.002 | 0.004 | 0.018 | 0.018 | 0.003 | 0.011 | 0.046 | 0.024 | 0.006 | 0.053 | 0.542 | 0.000 | 0.059 | 0.192 | 0.018 |
| Afro-Peruvians | 0.012 | 0.009 | 0.012 | 0.048 | 0.043 | 0.035 | 0.040 | 0.059 | 0.121 | 0.027 | 0.138 | 0.108 | 0.017 | 0.051 | 0.260 | 0.021 |
| Salvador | 0.005 | 0.007 | 0.014 | 0.067 | 0.052 | 0.008 | 0.000 | 0.012 | 0.124 | 0.009 | 0.199 | 0.379 | 0.065 | 0.015 | 0.039 | 0.005 |
| Bambui | 0.005 | 0.005 | 0.010 | 0.048 | 0.040 | 0.006 | 0.000 | 0.013 | 0.039 | 0.007 | 0.078 | 0.558 | 0.110 | 0.020 | 0.053 | 0.006 |
| Pelotas | 0.005 | 0.011 | 0.016 | 0.075 | 0.063 | 0.007 | 0.002 | 0.010 | 0.020 | 0.018 | 0.103 | 0.461 | 0.122 | 0.022 | 0.059 | 0.007 |

*The Kwa/Gur dataset includes approximately 35 ethno-linguistic groups, predominantly from the Kwa and Gur Niger-Congo linguistic group[6].
**The Nilotic dataset includes predominantly three ethno-linguistic groups in Northern Uganda (Langi, Acholi and Lugbara) from the Nilotic linguistic group[6].

### Extended Data Table 3. (Continuation)

| | African Proportion | RAP_Sandawe | RAP_Tswana | RAP_Herero | RAP_Mbukushu | RAP_LWK | RAP_Nilotics | RAP_MSL | RAP_GWD | RAP_Kwa/Gur | RAP_ESN | RAP_YRI |
|---|---|---|---|---|---|---|---|---|---|---|---|---|
| ACB | 0.885 | 0.001 | 0.004 | 0.003 | 0.017 | 0.034 | 0.002 | 0.067 | 0.030 | 0.380 | 0.071 | 0.390 |
| PUR | 0.189 | 0.022 | 0.009 | 0.016 | 0.092 | 0.104 | 0.013 | 0.063 | 0.283 | 0.110 | 0.072 | 0.215 |
| PLCO | 0.883 | 0.006 | 0.012 | 0.012 | 0.059 | 0.066 | 0.020 | 0.090 | 0.098 | 0.192 | 0.111 | 0.334 |
| ASW | 0.773 | 0.004 | 0.011 | 0.012 | 0.053 | 0.080 | 0.010 | 0.099 | 0.105 | 0.154 | 0.108 | 0.364 |
| CLM | 0.189 | 0.013 | 0.011 | 0.020 | 0.098 | 0.097 | 0.015 | 0.057 | 0.246 | 0.126 | 0.034 | 0.284 |
| Afro-Peruvians | 0.542 | 0.022 | 0.016 | 0.021 | 0.088 | 0.080 | 0.065 | 0.073 | 0.108 | 0.222 | 0.050 | 0.255 |
| Salvador | 0.497 | 0.009 | 0.013 | 0.028 | 0.134 | 0.105 | 0.017 | 0.000 | 0.024 | 0.249 | 0.019 | 0.401 |
| Bambui | 0.253 | 0.020 | 0.020 | 0.041 | 0.191 | 0.159 | 0.024 | 0.000 | 0.053 | 0.154 | 0.027 | 0.310 |
| Pelotas | 0.329 | 0.015 | 0.033 | 0.048 | 0.229 | 0.191 | 0.021 | 0.007 | 0.030 | 0.059 | 0.054 | 0.313 |

### Extended Data Table 4. GLOBETROTTER inferences of admixture events in the admixed American continent populations.

| Target admixed population[a] | Model[b] | Modeling estimates[c] | | | | | | Date of first potential Admixture event[d] | | | | Date of second potential Admixture event[d] | | | |
|---|---|---|---|---|---|---|---|---|---|---|---|---|---|---|---|
| | | p | $R^2$ | $FQ_1$ | $FQ_2$ | M | Events | Gen. | Date (CI) | 1st BSP (fcy) | 2nd BSP (fcy) | Gen. | Date (CI) | 1st BSP (fcy) | 2nd BSP (fcy) |
| PLCO, USA | Null | <0.01 | 1 | 1 | 1 | 0.29 | one date | 5.7 | 1749 (1749-1779) | CEU (0.13) | YRI (0.87) | - | - | - | - |
| ASW, USA | Null | <0.01 | 1 | 1 | 1 | 0.18 | one date | 6.4 | 1800 (1770-1800) | CEU (0.23) | YRI (0.77) | - | - | - | - |
| ACB, Barbados | Null | <0.01 | 1 | 0.943 | 1 | 0.22 | one date | 5.9 | 1800 (1800-1816) | CEU (0.12) | YRI (0.88) | - | - | - | - |
| PUR, Puerto Rico | Null | <0.01 | 0.998 | 0.994 | 1 | 0.52 | two dates | 4.9 | 1830 (1740-1920) | Ghana (0.11) | IBS (0.89) | 13.5 | 1560 (1500-1620) | Aymaras (0.24) | IBS (0.75) |
| CLM, Colombia | Null | <0.01 | 0.999 | 0.965 | 1 | 0.71 | two dates | 5.1 | 1830 (1710-1936) | Aymaras (0.28) | IBS (0.82) | 10.8 | 1650 (1574-1680) | Aymaras (0.34) | IBS (0.66) |
| Afro-Peruvians, Peru | Null | <0.01 | 0.999 | 0.999 | 1 | 0.54 | two dates | 5.1 | 1830 (1800-1950) | YRI (0.39) | Aymaras (0.61) | 8.3 | 1740 (1559-1770) | Aymaras (0.48) | IBS (0.52) |
| Salvador, Brazil | Null | <0.01 | 1 | 0.923 | 1 | 0.15 | one date | 8.2 | 1758 (1758-1758) | IBS (0.50) | YRI (0.50) | - | - | - | - |
| Bambuí, Brazil | Null | <0.01 | 1 | 0.988 | 0.999 | 0.98 | two dates | 4.1 | 1808 (1748-1838) | CEU (0.26) | IBS (0.74) | 8.6 | 1658 (1598-1658) | Ashaninkas (0.24) | IBS (0.76) |
| Pelotas, Brazil | Null | <0.01 | 0.999 | 0.964 | 1 | 0.94 | two dates | 4.8 | 1832 (1802-1892) | YRI (0.26) | IBS (0.74) | 8.5 | 1742 (1666-1772) | Ashaninkas (0.16) | IBS (0.84) |

The table shows the proportion and dates of inferred admixture events using GLOBETROTTER (Hellenthal, et al. 2014).
We defined the parental populations of our dataset (information on Table S1) as a donor and, each target admixed populations as a recipient.
(a) Target populations: CLM=Colombians from Medellin, PUR=Puerto Ricans from Puerto Rico, ACB=African Caribbeans from Barbados, PLCO= African Americans from East USA, ASW= Americans with African ancestry from South west USA.
(b)Two models were used, Main or traditional model and the Null-model which is more conservative including filters of spurious linkage disequilibrium involved in the admixture event.
(c)The modeling estimates were calculated using a 100 -bootstrap re-samplings; p=p-value, R2=goodness-of-fit of the tested model,
FQ1 = fit of a single admixture event, FQ2= fit of the first two principal components capturing the admixture events,
M=R2 of the model including a second date versus assuming only a single date of admixture (M>0.35 to infer multiple dates of admixture),
Events= inferred number of events for admixture process. (d)Inferred dates: Gen.= number of generations ago (we assume a generation time of 28 years);
Date (CI)= most likely year of the admixture event based on the mean birth year (Confidence Interval); BSP= Best Source Population,
Inferred best population sources for admixture event: IBS=Iberian population in Spain, YRI=Yoruba in Ibadan- Nigeria,
CEU=Utah Residents (CEPH) with Northern and Western Ancestry-USA, LWK= Luhya in Webuye, Kenya,
GHA=Ghana, AYM=Aymara population from Peru, showing in parenthesis the percentage of donation of each population source.

**Extended Data Table 5. Number of African individuals from ancestry-related origins that disembarked in the American continent regions and ports.**

| Destiny Proxies | Ports of Disembark | West-Central Africa1 | Southern/Eastern Africa1 | Western Africa1 |
|---|---|---|---|---|
| Northeast Brazil | "Bahia, port unspecified" | 801,153 | 474,593 | 7,089 |
| Southest Brazil | "Rio de Janeiro province" | 414,39 | 1,389,320 | 3,672 |
| United States | All US ports | 89,874 | 78,686 | 87,713 |
| Barbados | "Barbados, port unspecified" | 205,416 | 40,860 | 28,452 |
| Puerto Rico | "Puerto Rico, port unspecified" | 6,274 | 5,070 | 8,126 |
| Colombia | "Cartagena" | 22,058 | 91,875 | 94,522 |

1 Number of African Individuals disembarked in the destiny proxies of the slave trade that came from African regions related to the ancestral geographic origins as follows: "Gold Coast", "Bight of Benin" and "Bight of Biafra and Gulf of Guinea islands" = West-Central African origin; "West Central Africa and St. Helena" and "Southeast Africa and Indian Ocean islands" = Southern/Eastern African origin ; "Senegambia and offshore Atlantic" and "Sierra Leone" = Western African origin.

**Extended Data Table 6. The observed and expected proportions of genomic African ancestry clusters in the Americas based on demography and genetics.**

| Destiny Proxies | Genomic Data | WCA_Obs [1] | WCA_Exp [2] | SEA_Obs [1] | SEA_Exp [2] | WA_Obs [1] | WA_Exp [2] |
|---|---|---|---|---|---|---|---|
| Northeast Brazil | Salvador | 0.554 | 0.581 | 0.347 | 0.363 | 0.061 | 0.054 |
| Southest Brazil | Bambui and Pelotas | 0.330 | 0.202 | 0.508 | 0.768 | 0.096 | 0.028 |
| United States | PLCO and ASW | 0.537 | 0.431 | 0.233 | 0.291 | 0.196 | 0.277 |
| Barbados | ACB | 0.722 | 0.668 | 0.152 | 0.205 | 0.108 | 0.126 |
| Puerto Rico | PUR | 0.241 | 0.418 | 0.270 | 0.253 | 0.421 | 0.328 |
| Colombia | CLM | 0.347 | 0.282 | 0.323 | 0.372 | 0.274 | 0.345 |

1The current African ancestry clusters respect to the total African ancestry (see Table S2) inferred by the genomic data: West-Central African (WCA), Southern/Eastern African (SEA), Western African (WA).

2The weighted expected proportion of the ancestry cluster was calculated by the proportion of individuals that came from the ports (West-Central, Southern/Eastern and Western) related to the ancestral origins

(WCA, WA, SEA), respectively (Fig 1A, see details in the SI text and Extended Data Table 1 for the genomic proportions of ancestry used to obtain these results).

**Extended Data Table 7. Statistical comparison of the observed and expected distributions of the proportion of African ancestry clusters (relative to the total African ancestry).**

| Ancestry cluster | N | Observed mean | Observed inter-population variance | Adjusted theoretical distribution | Expected mean | P-Valor * |
|---|---|---|---|---|---|---|
| WCA in Southeastern Brazil | 1873 | 0.374 | 0.01641204 | Normal<br>Media = 0.202 SD = 0.1281095 | 0.202 | < 2.2e-16 |
| SEA in Southeastern Brazil | 1873 | 0.547 | 0.01331637 | Normal<br>Media = 0.768 SD = 0.1153966 | 0.768 | < 2.2e-16 |
| WCA in US | 55 | 0.547 | 0.002833085 | Weibull<br>Shape = 8.2295470 Scale = 0.3870523 | 0.365 | 3.33E-16 |
| WA in US | 55 | 0.191 | 0.001577122 | Normal<br>Media = 0.346 SD = 0.039713 | 0.346 | 3.33E-16 |
| WCA in the Americas | 3923 | 0.477 | 0.02534571 | Beta<br>$\alpha = 3.400755$ $\beta = 5.079897$ | 0.401 | < 2.2e-16 |

* p value of testing the null hypothesis that observed and expected distributions are not different, using the Kolmogorov-Smirnoff test. WCA = West-Central African;
WA = Western African; SEA = South/East Africa.
